# Transposons accelerate chromosomal speciation by centromere expansion and chromosome fission

**DOI:** 10.1101/2025.02.13.638213

**Authors:** Jui-Hung Tai, Tzi-Yuan Wang, Gwo-Chin Ma, Yu-Wei Wu, Tsung-Han Yu, Chen-Fan Wang, Te-Yu Liao, Shih-Pin Huang, Feng-Yu Wang, Yohey Terai, Tsung-Han Wu, Chun-Hua Hsu, Hon-Tsen Yu, Kwang-Tsao Shao, Shu-Miaw Chaw, Hurng-Yi Wang

**Author notes:** Corresponding authors. Email: KTS,; SMC,; HYW. These authors contribute equally to this work (Jui-Hung Tai, Tzi-Yuan Wang).

## Abstract

Chromosomal speciation poses a paradox: if rearrangements cause postmating isolation, how do they persist and spread? We propose that centromere remodeling, driven by transposable element accumulation, provides a mechanistic solution. In *Opsariichthys*, a hyper-diverse lineage of Asian river chub with 2N = 76–78 chromosomes, we identify ∼15 fission events coinciding with TE-fueled centromeric DNA expansion, potentially linked to a mutation in the PIWI1 protein. These expansions enable neokinetochore formation, allowing fissioned chromosomes to segregate faithfully, while subsequent rearrangements disrupt pairing and reduce gene flow. Comparative genomics reveals that fissioned chromosomes exhibit lower migration and greater divergence, acting as reproductive barriers. Our findings show that TE-mediated kinetochore duplication can facilitate extensive chromosome fission without meiotic disruption, providing a viable and previously underappreciated path to rapid speciation.

## Main Text

Chromosomal speciation model posits that fissions generate reproductive barriers that drive speciation (*1, 2*). In monocentric species, however, fission can produce acentric fragments or centromere imbalance, leading to meiotic failure (*3*). It remains unclear how chromosome fission occurs and contributes to reproductive isolation, despite the occasional occurrence of massive fission events in vertebrate evolution (*4*). One proposed solution is neokinetochore formation (*5, 6*): during gametogenesis, it can generate dicentric chromatids, and if these achieve bipolar attachment in meiosis, fission may yield two chromosomes with functional kinetochores and minimal phenotypic effect. Subsequent pericentric inversion may complicate pairing between fissioned and intact homologs, thus creating barriers to gene flow and driving speciation. While theoretically compelling, this model lacks strong empirical support (*7, 8*), and the mechanism triggering neokinetochore formation remains unknown.

Transposable elements (TEs) have recurrently shaped genomic variation and host diversity throughout evolution (*9*). Their activity drives chromosomal rearrangements— including fusions, fissions, translocations, and inversions—that can fuel evolutionary innovation and speciation (*10, 11*). We propose that active TE accumulation triggers mutagenic processes such as unequal exchange and replication slippage in the centromere (*12*), promoting the expansion of centromeric DNA and initiating neokinetochore formation.

The Opsariichthyini tribe (Asian river chub) (Cypriniformes: Xenocyprididae) includes small riverine minnows broadly distributed across East Asia, from Russia to northern Vietnam (*13*). This monophyletic group contains multiple genera, ranging from early- diverging lineages such as *Parazacco, Candidia*, and *Nipponocypris* to more recently diverged genera such as *Zacco* and *Opsariichthys*. Most species in the tribe have 2N = 48 chromosomes, except *Opsariichthys*, which exhibits a strikingly elevated count of 2N = 74 - 78 (*14, 15*). Interestingly, hybrids between *Zacco* and *Opsariichthys*, as well as between different *Opsariichthys* species, have occasionally been seen in natural environments where they are sympatric (*16, 17*). Despite the increase in chromosome number, overall DNA content remains conserved, implicating extensive chromosome fission rather than genome expansion as the primary driver of karyotypic change (*18*). Notably, *Opsariichthys* contains over 15 recognized species (*19*), markedly more than the typical three to four species found in other genera in the tribe (*20*). This exceptional diversity, alongside its tolerance to widespread fission, makes *Opsariichthys* a compelling system for testing chromosomal speciation models.

To investigate the relationship between chromosome fission and TE activity in *Opsariichthys*, and to assess whether fission contributes to accelerated speciation, we sequenced and annotated the genomes of *Candidia barbatus* (Cb), *Zacco platypus* (Zp), *O. pachycephalus* (Op), and *O. evolans* (Oe), each sampled from multiple populations. In parallel, we analyzed the karyotypes of these species to enable integrated analysis of chromosomal variation and fission patterns. For comparison, the genome and karyotype of *O. bidens* (Ob) were included in subsequent analyses based on published data (*21*). These five species capture the genomic and ecological diversity within the Opsariichthyini tribe. Our results suggest that chromosome fissions were initiated by lineage-specific TE activity, likely beginning in the common ancestor of Opsariichthyini. Fissioned chromosomes showed reduced migration and increased divergence relative to intact chromosomes, indicating their role in impeding gene flow and facilitating speciation.

### Karyotyping, genome assembly and annotation

Karyotypes were determined by examining multiple individuals from each species. No heteromorphic elements were detected between males and females. Cb and Zp both possess 2N = 48 chromosomes, while Op and Oe have 2N = 76 and 78, respectively (Fig. 1, fig. S1 and table S1). Despite the wide variation in chromosome number, the fundamental numbers are comparable across species. Genome assembly details are given in Supplementary Text (Genome assembly statistics; see also fig. S2, table S2-S6). Chromosome-level assemblies are consistent with karyotyping results and exhibit high completeness, with BUSCO scores exceeding 96% for all species (table S4). Transposable elements (TEs) account for 51% to 55% of the opsariichthyin genomes. Notably, the Opsariichthyini show a higher proportion of long interspersed nuclear elements (LINEs), particularly L2, and a significant increase in Gypsy within their long terminal repeats (LTRs) content compared to other cyprinids (table S5).

**Fig. 1.**
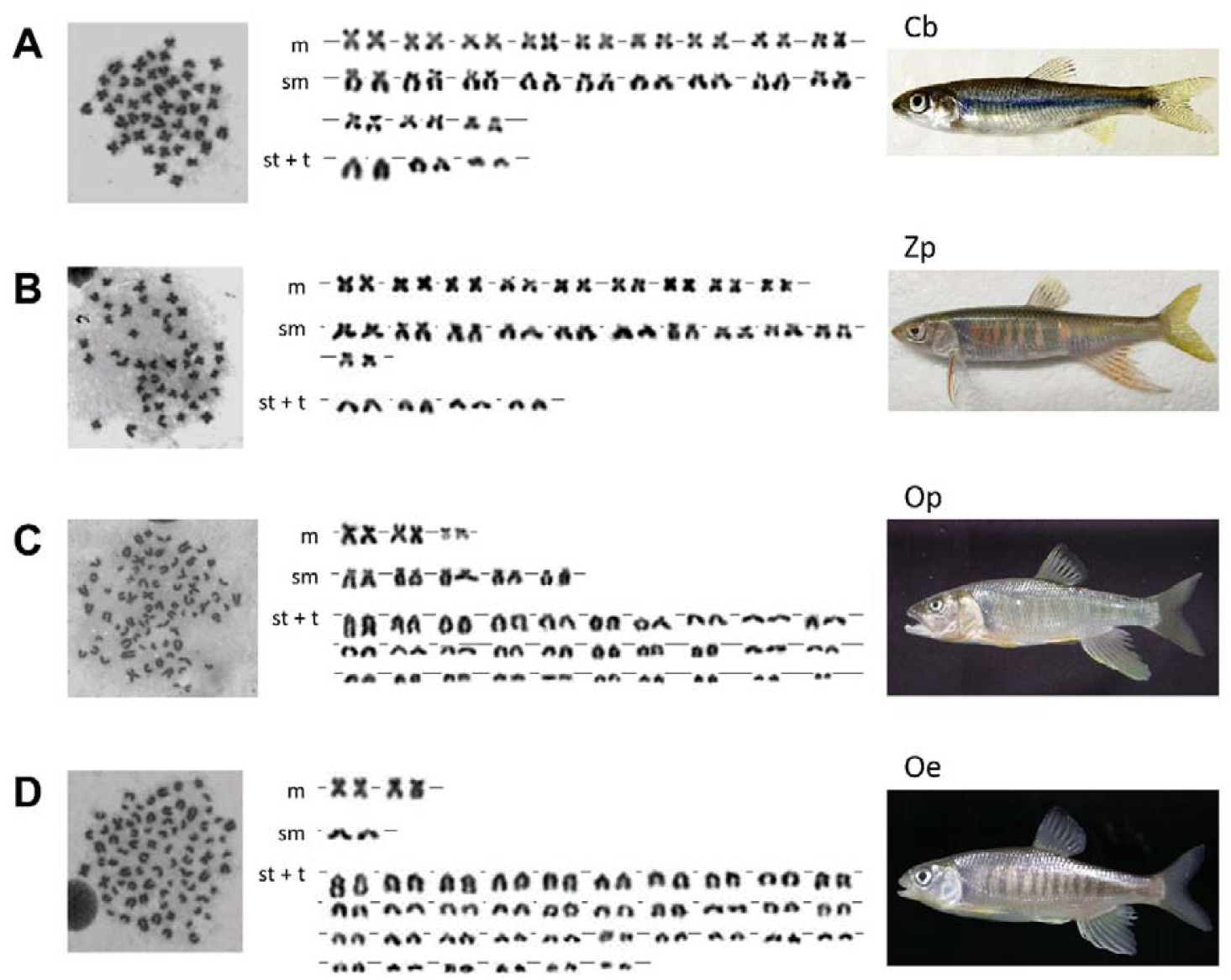
Karyotypes of opsariichthyins examined. **A**, Cb (*Candidia barbatus*) **B**, Zp (*Zacco platypus*) **C**, Op (*Opsariichthys pachycephalus*) and **D**, Oe (*O. evolans*). m: metacentric; sm: submetacentric; st: sub-telocentric; t: telocentric.

### Chromosomal evolution is associated with recent TE activities

Synteny analysis revealed that chromosome numbers in *Opsariichthys* increased dramatically from N = 24 to 38 or 39, driven by 14 to 15 chromosome fission events (Fig. 2). We identified ‘breaking regions’ in Cb and Zp, where segments of ancestral chromosome no longer align with their fissioned homologs (fig. S3), likely due to structural changes or sequence loss during fission. These breaking regions are enriched with Gypsy and L2, with Zp showing a higher Gypsy density than Cb (fig. S4A). This enrichment extends to the homologous regions in the fissioned chromosomes of Op, Oe, and Ob (fig. S3).

**Fig. 2.**
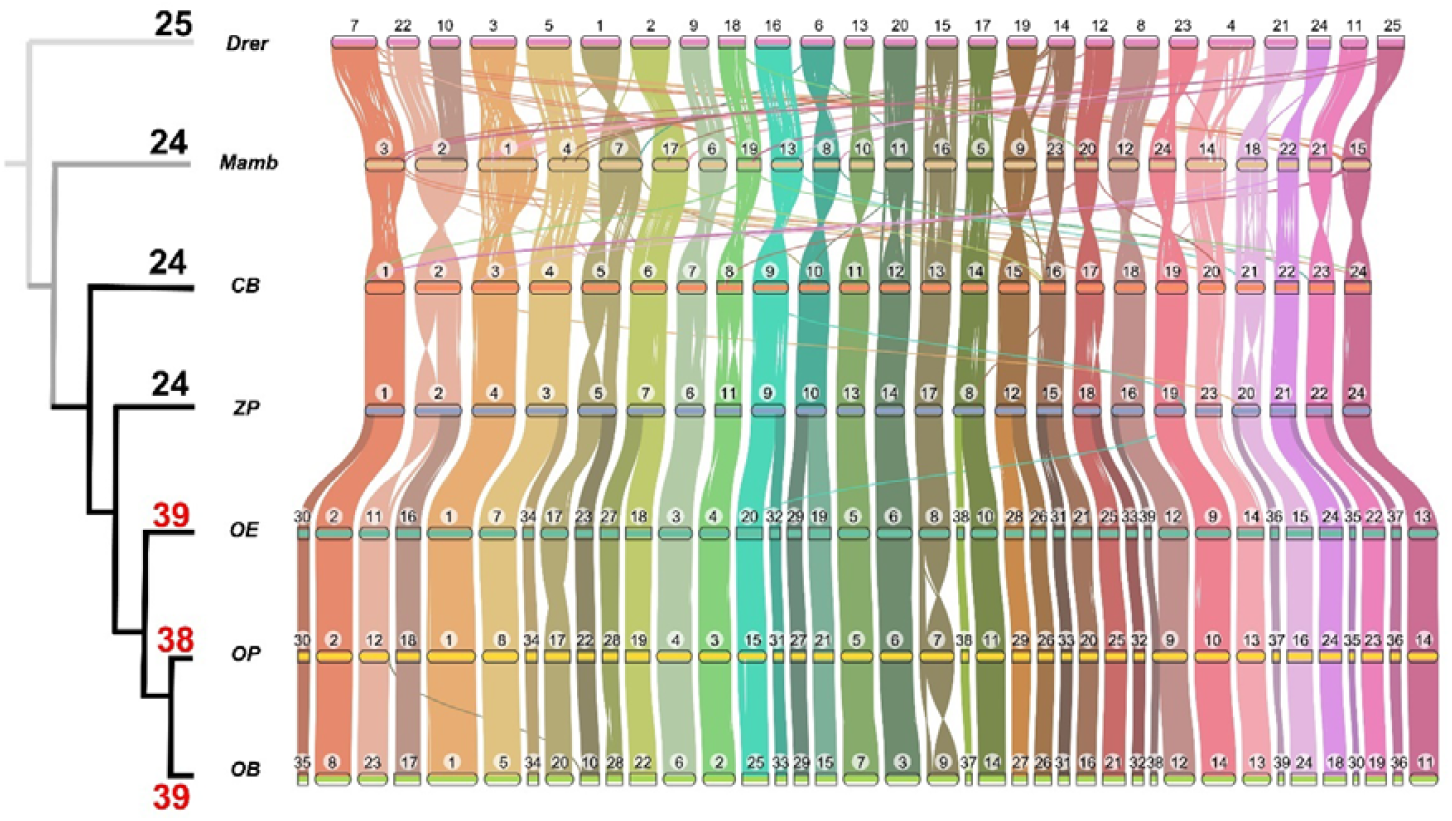
The phylogeny (left panel) and syntenic relationships (right panel) among Opsariichthyini tribe species and the outgroups. Dark black branches represent lineages of the Opsariichthyini tribe. The numbers represent the chromosome number (N). Drer: *Danio rerio*; Mamb: *Megalobrama amblycephala*; Cb: *Candidia barbatus*; Zp: *Zacco platypus*; Oe: *Opsariichthys evolans*; Op: *O. pachycephalus*; Ob: *O. bidens*

Repeat landscapes revealed recent expansions of L2 and Gypsy (k-distance < 5%; young TEs) across all examined opsariichthyin genomes (fig. S4B). In Cb and Zp, 56% and 43% of L2 insertions, respectively, are recent, increasing to 75% and 72% within breaking regions (fig. S4C). Recent Gypsy expansion is more pronounced in Zp, increasing from 17% genome-wide to 31% in breaking regions, whereas in Cb, only 10% of Gypsy are recently inserted in both genomic and breaking regions. Furthermore, 14 of the 15 breaking regions in Cb show recent L2 expansion and 8 show recent Gypsy expansion (table S7; fig. S5), while all breaking regions in Zp exhibit recent expansion of both elements (table S8; fig. S6). These observations suggest that L2 and Gypsy expansion occurred asynchronously, with L2 expanding before Gypsy.

### Chromosome fission within centromere regions

To investigate the mechanism of chromosome fission, we identified putative centromere regions by searching for regions containing centromeric DNA, low recombination rates, and enriched with L2 and Gypsy (Supplementary Text: Centromere identification; fig. S5-S10). In general, karyotyping results derived *in silico* analysis and microscopic observations are largely consistent (Fig. 1 and table S1), demonstrating that the putative centromeres identified are authentic. In Cb, 10 of 15 breaking regions overlapped with putative centromeres (fig. S5; table S7), while in Zp, all 15 coincided (fig. S6; table S8). The difference is primarily due to chromosomal rearrangements, especially pericentric inversions, that occurred during evolution (fig. S11). Recent L2 expansions were observed in centromeres across all species. Gypsy expansions were prevalent in nearly all centromeres of Zp, Op, Oe, and Ob, but occurred in only 14 of Cb’s 24 centromeres (table S7-S11). These patterns, consistent with breaking regions, support earlier L2 expansion than Gypsy.

Notably, in chromosomes that later underwent fission in *Opsariichthys*, the corresponding regions in Zp already exhibit significantly greater centromeric DNA content and broader centromeric spans than in Cb (Fig. 3A). While the number of L2 in the centromere regions is similar among species, that of Gypsy is significantly greater in Zp (Fig. 3B). These findings suggest that recent Gypsy expansion may have promoted both centromeric DNA duplication and centromere expansion through unequal exchange and replication slippage. Together, these processes could facilitate neokinetochore formation, leading to dicentric chromosomes (*6*), in which each fissioned product retains a functional kinetochore for proper segregation.

**Fig. 3.**
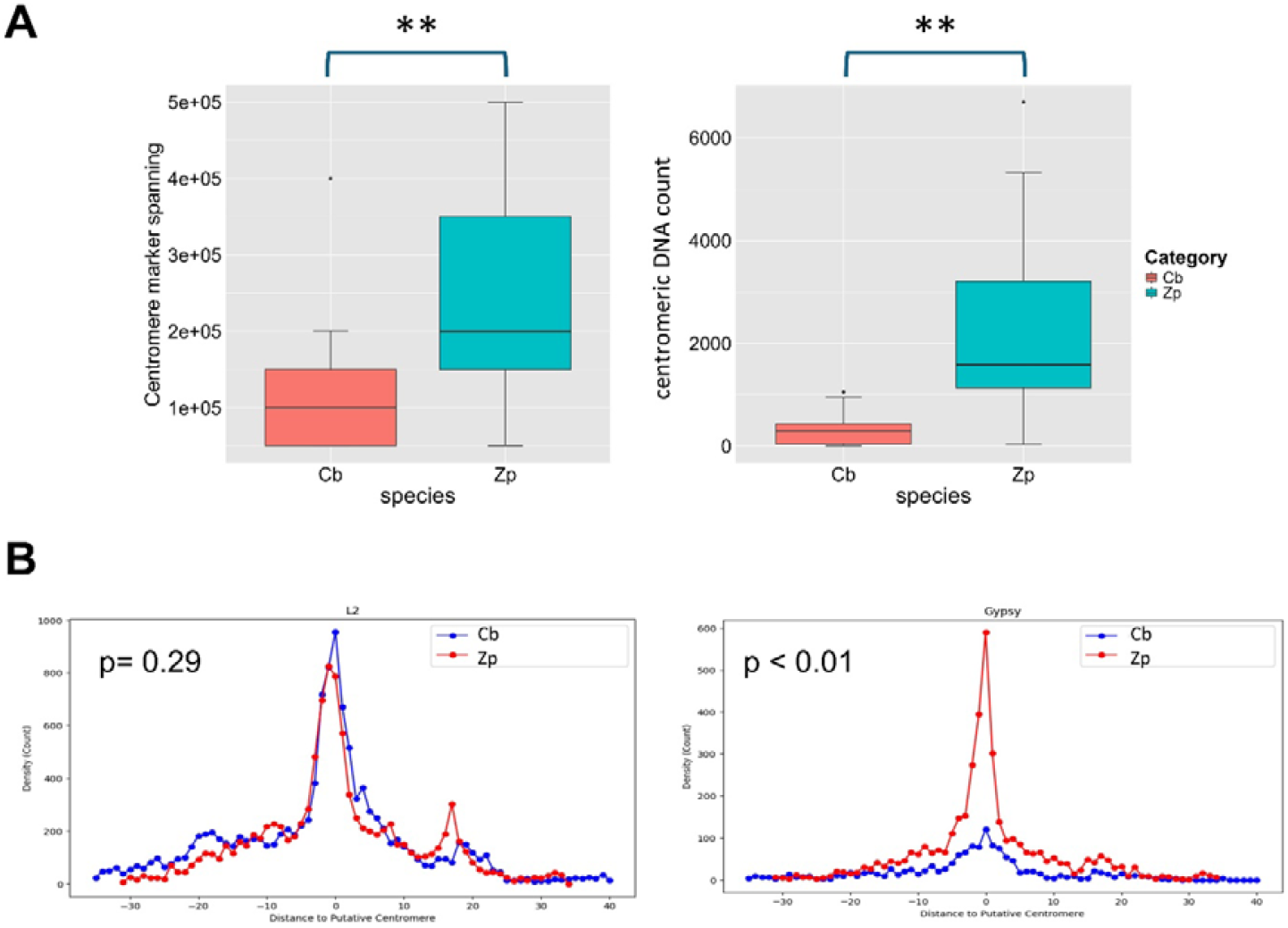
The distribution of centromeric DNA, L2, and Gypsy within putative centromeres of Cb and Zp in fissioned chromosomes. **A**, The left panel displays the regions spanned by centromere markers, and the right panel shows their counts in fissioned chromosomes. **B**, The distribution of L2 (left panel) and Gypsy (right panel) around centromeres in fissioned chromosomes. X-axis represents distance to the center of putative centromere regions. ** p < 0.01

### Insertion preferences and interaction between L2 and Gypsy

The expansion of young L2 in putative centromeres suggests that the reduced recombination rates in these regions limit transposon removal, enabling their accumulation over time (*22*). This is evidenced by a negative correlation between young L2 density and recombination rates in Zp (r = −0.44) and Cb (r = −0.25) (fig. S12A). Because centromeres can shift over time (fig. S11), ancient L2 (k-distance > 5%) show reduced or no correlation with recombination rate (Zp: r = –0.11; Cb: n.s.) (fig. S12B).

The expansion of L2 preceding that of Gypsy suggests a potential interdependence (fig. S13), with expansion of Gypsy possibly relying on a genomic context shaped by L2. LTR insertion is typically marked by a 5-bp target site duplication (TSD) flanked by palindromic motifs generated during integration (*23*). Using LTR_finder and LTR_retriever, we identified a conserved palindromic motif flanking the TSD in opsariichthyin species: TNACA-TSD- TGTNA (fig. S14A). Similar motifs are present in L2 (fig. S14B), suggesting that each L2 replication and transposition event creates a new target site for subsequent Gypsy integration.

### Chromosome fission decrease gene flow and increase genetic divergence

According to the chromosome speciation model, chromosomal rearrangements can lead to the formation of a reproductive barrier that promotes speciation. Divergence at loci on rearranged chromosomes is also expected to exceed that on non-rearranged chromosomes. To test this, we used gIMble (*24*) to calculate the blockwise site frequency spectrum (bSFS) between Zp and Oe, as well as between Zp and Op. The isolation with migration (IM) model was the best fit (table S12). Migration near the breaking regions was notably reduced (Fig. 4A and fig. S15-S16), and fission chromosomes exhibited significantly lower migration rates than non-fission chromosomes in both comparisons (p < 0.05) (Fig. 4B).

**Fig. 4.**
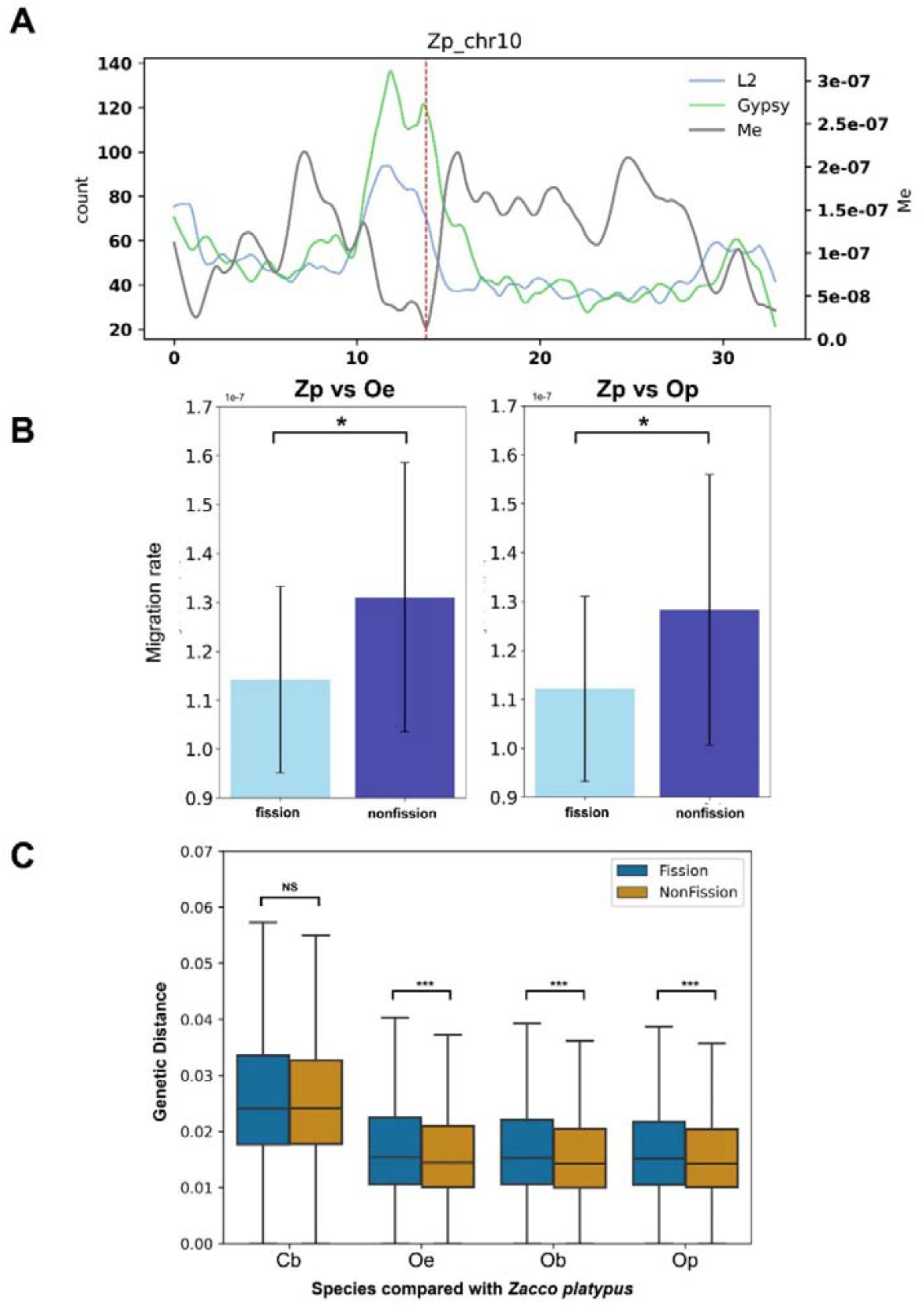
Gene flow was reduced in fissioned chromosomes. **A**, Migration rate (Me) between Zp (*Z. platypus*) and Oe (*Opsariichthys evolans*) across various chromosomes, exemplified here by chromosome 10 of Zp, with additional data for other chromosomes presented in fig. S8-S9. Red dash line indicates breaking regions. **B**, The average Me estimates for fission and non-fission chromosomes between Zp and Oe (left panel) and between Zp and Op (right panel). **C**, Average genetic distance of single copy genes on fission and un-fissioned chromosomes between Zp and each of Cb, Oe, Ob, and Op. * p < 0.05; *** p < 10-6, NS: not significant.

We next calculated the genetic divergence using single-copy genes. Chromosomes that underwent fission in *Opsariichthys* showed significantly greater genetic distances than non-fission chromosomes in all comparisons with Zp (p < 10^−6^) (Fig. 4C). By contrast, when comparing Cb and Zp, where both sets of chromosomes retained the ancestral, unfissioned state, no difference in divergence was detected between the two chromosome sets.

It is generally believed that fission does not reduce recombination. Indeed, fissioned chromosomes in both Op and Oe exhibited higher average recombination rates than non- fissioned chromosomes (fig. S17). In Zp, the homologous chromosomes showed comparable recombination rates across both groups.

### Mutations in PIWI1 protein associates with L2 expansion and species diversification

While the recent Gypsy expansion occurs in multiple cyprinids (fig. S18A), recent L2 expansion appears unique to the Opsariichthyini, suggesting possible shifts in transposon silencing. Given the critical role of PIWI1 in transposon repression (*25*), we compared PIWI1 sequences between opsariichthyins and other cyprinids (Supplementary Text: Comparative Analysis of Piwi1 Protein). All opsariichthyins share two unique amino acid substitutions, G348S and V380I, in the PAZ domain, a region essential for piRNA binding and TE silencing (fig. S18C). V380I is particularly notable, as it lies adjacent to the nucleic acid-binding interface at residue 379 (*26*).

Repeat landscape analysis shows that cyprinids with the ancestral V380 lack L2 expansion (fig. S18A), while *Triplophysa* (Nemacheilidae), a cyprinid outgroup carrying V380I, has undergone recent L2 expansion (fig. S18B). Structural modeling using AlphaFold 3 revealed that residue 380 lies within the piRNA binding pocket of the PAZ domain (fig. S19A). Although it does not directly contact the binding interface (fig. S19B, upper panel), substitution of valine with the bulkier and more hydrophobic isoleucine (V380I) may subtly alter local conformation (fig. S19B, lower panel), potentially impairing piRNA loading or destabilizing the complex (*27, 28*). Given the phylogenetic distance between Opsariichthyini and *Triplophysa*, the parallel emergence of both the V380I mutation and L2 expansion suggests convergent evolution. These observations point to a shared mechanism, whereby V380I interferes with PIWI1 function, diminishing transposon silencing and enabling L2 proliferation.

In *Triplophysa*, chromosome numbers remain stable (2N = 48–50), yet karyotypes and fundamental numbers vary substantially (*29*), coinciding with L2 expansion (fig. S18). Notably, *Triplophysa* comprises 147 described species, far exceeding the family average of 16, suggesting a possible link between TE activity and elevated species diversification. Together, these findings implicate the V380I mutation in reshaping genome architecture and possibly contributing to lineage-specific diversification.

## Discussion

### Transposon-fueled centromere remodeling sets the stage for chromosome fission

Through comparative analysis of opsariichthyin genomes and interspecific gene flow, we provide direct evidence linking TE activity to chromosomal evolution and species diversification (Fig. 5A). L2 expansion is specific to Opsariichthyini within Cyprinoidei (Fig. 3B and fig. S18A) and is enriched in putative centromeres (Fig. 5B), where reduced recombination likely facilitates accumulation over time (*22*). L2 elements also harbor sequence motifs preferentially targeted by Gypsy, guiding its insertion into centromeric regions (fig. S14B). Moreover, LTRs frequently insert into the 3’ UTRs of other LTRs, especially Gypsy (*30*), creating a positive feedback loop that concentrates both elements in centromeres.

**Fig. 5.**
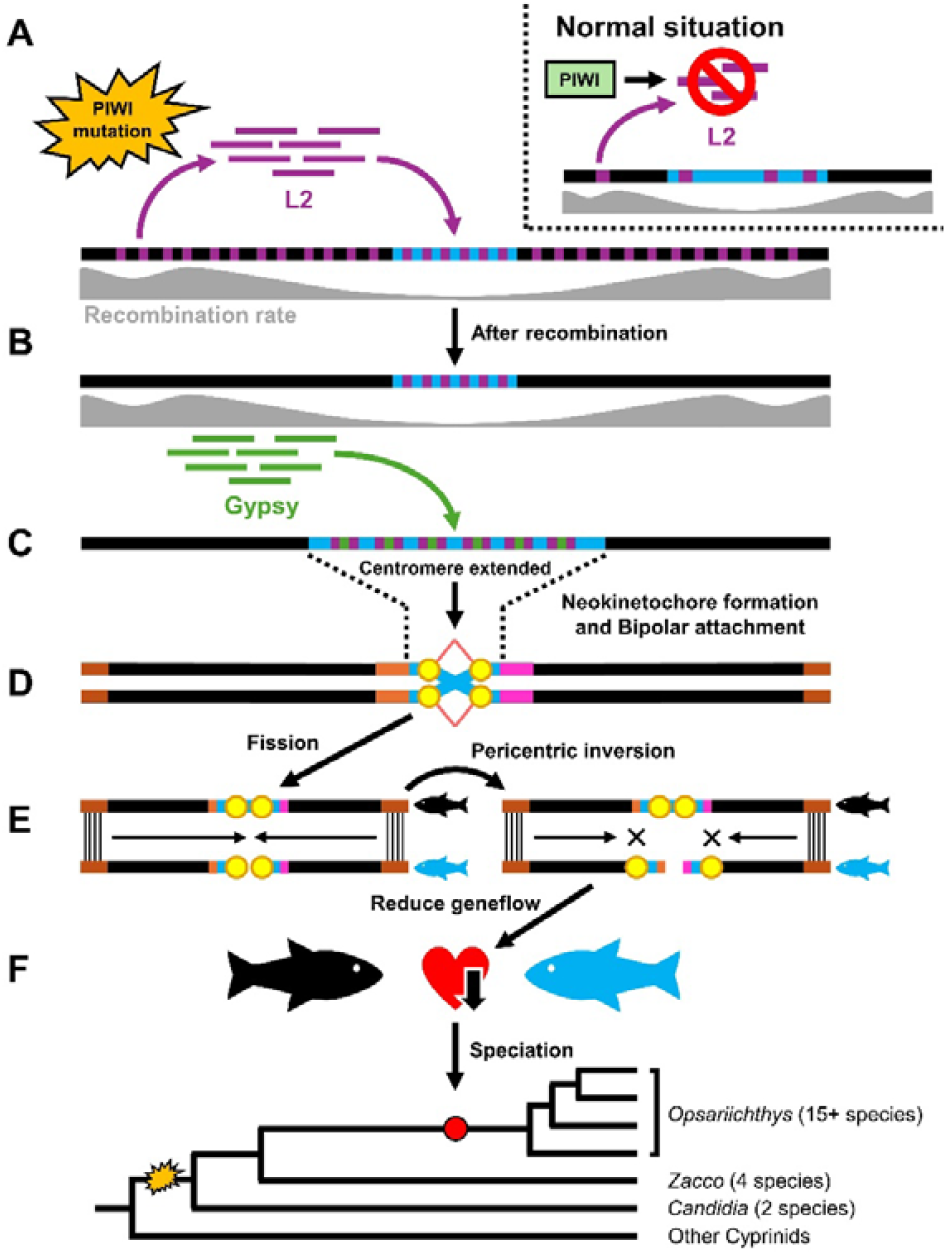
Transposable element expansion facilitates extensive chromosome fissions leading to species diversification. **A**, Under normal conditions (inset at the top right), the PIWI1 protein suppresses TE activity. However, mutations in the PAZ domain of the *piwi1* gene in the ancestor of the Opsariichthyini enabled the expansion of L2 within the genome. **B**, Reduced recombination within centromeres likely contributed to the accumulation of L2. **C**, L2 elements harbor sequence motifs preferentially for Gypsy, guiding its insertion into these motifs within centromeric regions and resulting in centromere expansion. **D**, Centromere expansion may facilitate neokinetochore formation, resulting in dicentric chromosomes. During replication, dicentric chromosomes with opposing bipolar attachment undergo chromosome fission. **E**, Following fission, the distal ends of the daughter chromosomes remain intact, ensuring proper pairing, while each of the two resulting chromosomes is equipped with a kinetochore, which enables accurate segregation (left). At later stage, however, chromosome rearrangement such as pericentric inversion may contribute to centromere relocation (right). **F**. Such rearrangements can act as reproductive barriers, reducing gene flow and driving speciation. Consequently, the species diversity of *Opsariichthys* increased.

Active TE accumulation may trigger mutagenic processes, such as unequal exchange and replication slippage in centromere(*12*), leading to further expansion of repetitive sequences including centromeric DNA. This is evident in Zp, where chromosomes that underwent fission after *Opsariichthys* diverged from *Zacco* contain significantly more centromeric DNA and exhibit centromere expansion compared to homologs in Cb (Fig. 5C), likely due to a recent Gypsy expansion (Fig. 3). According to kinetochore reproduction theory, expanded centromeric DNA increases the likelihood of neokinetochore formation (*5, 6*) (Fig. 5D). In the presence of kinetochores, dicentric chromosomes may arise. During replication, chromatids with monopolar spindle attachment segregate normally, while those with opposing bipolar attachment undergo fission (Fig. 5D and Fig 1B of ref 5).

Of the 15 identified breakpoints, 10 coincide with putative centromeres in Cb (fig. S5; table S7), while the remaining five, initially non-centromeric in Cb, became centromeres in Zp via pericentric inversions (fig. S11), resulting in all 15 regions perfectly match centromeres in Zp (fig. S6; table S8). Notably, similar associations between active TEs and chromosomal breakpoints have been observed in primate evolution (Supplementary Text: TEs in the breaking region of macaque chromosome 7).

### How fissioned chromosomes escape meiotic disruption and drive speciation

Following fission, accurate pairing of homologous chromosomes is required for proper segregation during meiosis. Centromeres have traditionally been considered essential not only for segregation but also for homologous pairing (*3*), raising concerns that new centromeres might disrupt this process. However, studies in wheat and *Arabidopsis* suggest centromere may not play a direct role in meiotic pairing (*31, 32*). Instead, chromosome pairing initiates at both ends and extends toward the center during early meiosis, indicating that pairing is end-driven rather than centromere-dependent (*33*). This provides a mechanistic explanation for how massive chromosome fission, as observed in *Opsariichthys*, can occur without disrupting meiotic pairing or segregation (Fig. 5E, left). Furthermore, selection may favor the retention of fissioned over intact homologs, especially if smaller chromosomes are more likely than larger ones to segregate without error during cell division (*7, 34*). As a result, fission karyotypes can rapidly increase in frequency within local populations without deleterious effects. For further evidence that frequent fission can proceed without deleterious effects in Opsariichthys, see Supplementary Text (Frequent fission and fusion in *Opsariichthys* genomes).

At later stages, additional rearrangement, such as pericentric inversion, common in the Opsariichthini (fig. S11), can relocate centromeres (Fig. 5E, right). Such relocation may interfere with pairing between fissioned and unfissioned homologs, thereby impeding gene flow, promoting genetic isolation, and ultimately facilitating speciation (Fig. 5F). Supporting this, gene flow between *Zacco* and *Opsariichthys* is reduced at chromosomal breakpoints, with fissioned chromosomes showing significantly lower migration estimates and greater genetic divergence than un-fissioned ones (Fig. 4).

Taken together, these findings invite a fresh perspective on the chromosomal speciation model, which traditionally posits that rearrangements such as fissions promote speciation by impeding gene flow, yet at the cost of meiotic dysfunction due to centromere imbalance. However, perhaps the paradox lies in the premise itself: the longstanding notion that chromosome rearrangements are inherently deleterious may overlook mechanisms that allow them to persist. Our results suggest that kinetochore duplication and end-driven pairing provide a viable path for fissioned chromosomes to segregate correctly and spread within populations, quietly laying the groundwork for reproductive isolation and speciation.

## Supporting information

Supplementary informatioms

## Acknowledgements

The authors wish to express their gratitude to Professor Chi-Chang Liu for providing valuable samples. Additionally, we thank the NGS High Throughput Genomics Core at the Biodiversity Research Center, Academia Sinica, for their support in library preparation and Hi-C sequencing. We are deeply grateful to Professor Shau-Ping Lin and Professor Isheng Jason Tsai for their invaluable advice throughout this research.

## Author contributions

Conceived the idea: TYW, KTS, SMC, HYW

Collected samples: JHT, TYW, GCM, THY, CFW, TYL, SPH, FYW, YT

Performed the experiment: TYW, GCM, THW, CHH, HTY, YT

Performed the analysis: JHT, TYW, CFW, YWW, THY

Supervised the project and secured the fundings: KTS, SMC, HYW

Wrote the manuscript: JHT, TYW, THY, KTS, SMC, HYW

## Data and materials availability

All data in this study has been deposited to NCBI bioproject under accession PRJNA1159521 (Cb), PRJNA1160456 (Zp), PRJNA1192024 (Op), PRJNA1192752 (Oe) and PRJNA1203011 (Hybrids). The details were listed in Supplementary Tables and Supplementary Information.

## Supplementary Materials

Materials and Methods

Supplementary Text

Figs. S1 to S21

Tables S1 to S13

References (35–85)

