## Supplementary informatioms for "Transposons accelerate chromosomal speciation by centromere expansion and chromosome fission"

**The PDF file includes:**

Materials and Methods

Supplementary Text

Figs. S1 to S21

Tables S1 to S13

References

**Materials and Methods**

Chromosome Analysis

Species, including *Candidia barbatus* (Cb), *Zacco platypus* (Zb), *Opsariichthys evolans* (Oe) and *Opsariichthys pachycephalus* (Op), used for Analysis are listed in Table S1 (See also Fig. S19). Approximately 16 hours before the experiment, a yeast solution (0.05g yeast powder, 0.05g glucose, 2ml ddH2O) was injected into the dorsal musculature. One hour before euthanasia, 0.5% colchicine in 0.8% NaCl solution (8 μL per 1 g of body weight) was administered intraperitoneally. Tissues were minced using a glass tissue grinder, mixed with 3ml of 0.56% KCl, and gently resuspended. The suspension was filtered through cheesecloth to remove debris and transferred to a 15ml tube. After swelling in the hypotonic solution for 30 minutes, the suspension was centrifuged at 1600 rpm for 5 minutes. The supernatant was discarded, and the pellet was resuspended in residual solution. Gradually, 5ml of fixative (ethanol: glacial acetic acid = 3:1) was added. After 20 minutes at room temperature and subsequent centrifugation, the cells were resuspended in fixative. This fixation and centrifugation cycle was repeated once more. Finally, the supernatant was discarded, leaving 1-0.5ml of fixative. To prepare slides, the cell suspension was pipetted onto slides from a height of 20-25 cm above a slide on an 80°C heat slab. Slides were then either heated at 80°C for 2-3 hours or left at room temperature overnight. Slides were treated with Trypsin-EDTA at 37°C for one minute, halted with 95% ethanol, rinsed with ddH2O, and stained with Wright Dye for 1-3 minutes. After washing with tap water, slides were dried and stained with 6% Giemsa solution for 5-10 minutes, then rinsed and examined. Chromosomes were classified into three types, metacentric, submetacentric, and sub-/telocentric-like, following the classification method of Levan, Fredga and Sandberg (*35*).

Sample Collection

Species, including *Candidia barbatus* (Cb), *Zacco platypus* (Zb), *Opsariichthys evolans* (Oe) and *Opsariichthys pachycephalus* (Op), used for genome sequencing and assembly are listed in Table S2 (See also Fig. S19). Tissues for RNA sequencing were dissected from fish body and preserved in RNAlater™ Stabilization Solution (Cat: AM7021, Invitrogen, Thermo Fisher Scientific Inc.) at -80°C freezer. The other tissues were preserved in 100% ethanol at -80°C freezer. All experiments in this study were performed in accordance with guidelines of the animal ethics committee and were approved by the Institutional Animal Care and Use Committee (IACUC 15-12-923), Academia Sinica.

DNA and RNA sequencing

Approximately 30-50 mg of muscle tissue was processed using proteinase K and Buffer G2, which includes RNAse A, from the Genomic DNA Buffer Set (Cat: 19060, Qiagen). DNA extraction was performed using the Genomic-tip 20/G (Cat: 10223, Qiagen) following the manufacturer’s instructions.

For Nanopore sequencing, 5-8μg of genomic DNA was prepared by end-repair and ligation using the KAPA Hyper Prep Kit (Cat#KR0961, Kapa Biosystems, Wilmington, MA, USA) according to the supplier's guidelines. The prepared DNA library was combined with LB and SQB buffer from the Nanopore Ligation Sequencing Kit (SQK-LSK109, Oxford Nanopore Technologies, UK), loaded into a flow cell (R9.4.1; FLO-MIN106), and sequenced using MinION devices over a period of 24-72 hours.

For Illumina sequencing, genomic DNA was fragmented with a Covaris 2 device and its concentration was determined using a Qubit 2.0 Fluorometer. A sequencing library for short reads was generated from 200 ng of fragmented DNA utilizing the TruSeq Nano DNA High Throughput Library Prep Kit (Cat. No.20015965) and subsequently sequenced on an Illumina X-ten by GENOMICS BIOSCIENCE & TECHNOLOGY CO., LTD, Taipei, Taiwan.

For RNA sequencing, tissues (5 to 30 mg) were dissected and homogenized in 600 µl of Trizol for RNA extraction. RNA was then isolated using the RNeasy® Plus Mini Kit (Cat: 74134, Qiagen), and its quality was assessed with the Agilent 2100 Bioanalyzer. A sequencing library for short reads was created from about 1.5μg of RNA with the TruSeq Stranded mRNA Library Prep Kit (Cat. No. 20020595) and sequenced on the Illumina NovaSeq 6000 by the services of GENOMICS.

Genome Assembly and Size Estimation

In this study, we assembled four genomes, Cb, Zp, Oe, and Op. The nanopore long reads were assembled using Flye v2.9.2-b1786 (*36*) with “-m 10000 -meta” parameter. The resulting assemblies were corrected using POLCA (*37*) on Illumina short reads.

To assemble a chromosome-level genome, we employed scaffolding techniques using HiC and HiRise. Samples were prepared following the protocol of Lieberman-Aiden, van Berkum, Williams, Imakaev, Ragoczy, Telling, Amit, Lajoie, Sabo, Dorschner, Sandstrom, Bernstein, Bender, Groudine, Gnirke, Stamatoyannopoulos, Mirny, Lander and Dekker (*38*). In brief, chromatin was fixed using formaldehyde. The fixed chromatin was digested with DpnII, the 5’ overhangs filled in with biotinylated nucleotides, and the blunt ends were subsequently ligated. After ligation, the crosslinks were removed and DNA was purified. Any unligated biotin was eliminated from the purified DNA, which was then fragmented to approximately 350 bp. Sequencing libraries were generated with NEBNext Ultra enzymes and Illumina-compatible adapters. Biotin-containing fragments were isolated using streptavidin beads prior to PCR enrichment for each library. These libraries were then sequenced on an Illumina HiSeq X platform. After sequencing, Dovetail HiC library reads were processed with Juicer (*39*) to generate high-resolution contact maps. These maps were then served as the input file for 3d-dna (*40*) to anchor the address of contigs. The final visualization of the anchored contigs was conducted using Juicebox (*41*).

Genome size was estimated by Jellyfish (*42*) with parameter *k=17*. For estimating genome completeness, we blasted with “Actinopterygii” ortholog set, which consists of 3640 core genes, by using Benchmarking Universal Single-Copy Orthologs v5 (BUSCO v5) (*43*). The corrected nanopore reads were mapped back to the assembled genome to calculate the mapping rate using BWA v0.7.17-r1188 (*44*).

Transposable Elements Annotation

Transposable elements (TEs) in the genome were identified using both *de novo* prediction and homology search methods. For *de novo* prediction, we primarily used the repeat library retrieved from RepeatModeler v2.0.2 (*45*) with default parameters, supplementing it with LTR_FINDER_parallel v1.2 (*46*), LTR_FINDER v1.07 (*47*) and LTR_retriever v2.9.0 (*48*). For the homology search, we initially used the script famdb.py from RepeatMasker v4.1.2-p1 (*49*) to obtain “Cyprinidae” repetitive sequences from Dfam (ver. 3.5, 22-Dec-2021) (*50*) and Repbase (ver. 20181026) (*51*). Next, we downloaded repetitive sequences of *Ctenopharyngodon idellus* and *Danio rerio* from FISHTEDB (<http://www.fishtedb.org/>) (*52*). After concatenating these libraries, RepeatMasker was used for genome masking. We also utilize the script *calcDivergenceFromAlign.pl* to calculate the Kimura distance for each TEs family and used *createRepeatLandscape.pl* to graph the repeat landscape. To plot the distribution and proportion of TEs across chromosomes, a custom Python script was used with a window size of 50k bp.

Gene Annotation

Braker2 (*53*) was used for the prediction of protein-coding genes. Protein-based and RNA-based prediction were predicted separately. Protein sequences from various fish species, including *Astyanax mexicanus*, *Carassius auratus*, *Chanos chanos*, *Clupea harengus*, *Danio rerio*, *Paramormyrops kingsleyae*, *Salmo salar* and *Scophthalmus maximus*, were downloaded from GenBank (Supplementary Text: Published genome used in the analysis). RNA sequences were derived from sequencing of tissues detailed in Table S2. In order to annotate the published genome of Ob (NGDC, GWHBEIO00000000), the RNA sequences of Ob were downloaded from NCBI (SRR15969039). The integration of predictions was facilitated by TSEBRA (*54*), using the 'pref_braker1.cfg' configuration to prioritize RNA-based predictions. Subsequent filtering of preliminary gene predictions involved two criteria: 1) incompleteness of the prediction, identified by the absence of an initial methionine, a missing stop codon, or the presence of multiple stop codons within the sequence; and 2) sequences shorter than thirty-three amino acids.

Published Genomes Used in the Analysis

We analyzed the genomes of ten cyprinid species and two Nemacheilidae species as an outgroup for comparative genome analysis, transposable element analysis, or Piwi1 protein sequence comparison. The cyprinid species included *Danio rerio*(Drer) (*55*), *Danionella cerebrum*(Dcer) (*56*), *Labeo rohita*(Lroh) (*57*), *Sinocyclocheilus grahami*(Sgra) (*58*), *Onychostoma macrolepis*(Omac) (*59*), *Puntigrus tetrazona*(Ptet)(GCF_018831695.1), *Pimephales promelas*(Ppro) (*60*), *Anabarilius grahami*(Agra) (*61*), *Megalobrama amblycephala*(Mamb) (*62*) and *Opsariichthys bidens*(Obid) (*21*). Two Nemacheilidae species included *Triplophysa tibetana*(Ttib) (*63*) and *Triplophysa bleekeri*(Tble) (*64*). The details of the aforementioned genome statistics are provided in Table S4.

Synteny analysis among species

Synteny analysis was conducted by MCscan (*65*) and Synima (*66*). MCscan was used to visualize synteny among species. The fine scale synteny blocks were retrieved from the results of Synima and were further used to identify breaking regions. The breaking regions are the segment of the original chromosome that cannot be readily aligned with the sequences in two fissioned homologous chromosomes. These regions represent genomic areas where disruptions in synteny occur, likely due to structural changes or the loss of genetic material during the chromosome fission process.

Candidate centromere sequences

The procedures for identifying centromere markers was based on the approach of Melters, Bradnam, Young, Telis, May, Ruby, Sebra, Peluso, Eid, Rank, Garcia, DeRisi, Smith, Tobias, Ross-Ibarra, Korf and Chan (*67*). In summary, the short read assembler PRICE (*68*) was utilized to assemble reads from the dataset previously used for genome polishing. The script provided by Melters, Bradnam, Young, Telis, May, Ruby, Sebra, Peluso, Eid, Rank, Garcia, DeRisi, Smith, Tobias, Ross-Ibarra, Korf and Chan (*67*) was then used to identify high order repeats, which were considered likely candidates for centromere repeats due to their high abundance as tandem repeats in the genome. Due to variations in the length of candidate centromere sequences, they were aligned and visualized using the AliView software (*69*) to discern the simplest repeat units, with manual adjustments made to the sequences as necessary. These procedures were performed for each species, revealing that the identified centromere sequences were fundamentally similar across all species examined. Subsequently, the putative centromere sequences were blasted against their respective genomes to determine their positions and copy numbers on chromosomes.

Recombination rate estimation

Illumina short reads were mapped to the reference genome using NVIDIA Parabricks v4.3.1 (fq2bam) (*70*). Duplicate reads were removed using samtools rmdup (*71*). Bcftools mpileup (-q 20) and call (-m) (*71*) was used to generate VCF files for each individual. These files were filtered using bcftools view to retain sites with depths either greater than five or less than the mean depth plus two standard deviations. Subsequently, VCF files from the same species were merged using bcftools merge, and variants with data missing in fewer than two individuals were kept using bcftools view (-i 'N_MISSING<2'). The VCF files were phased using using Beagle v5.4 (*72*). Recombination rate estimation was carried out with LDhelmet v1.9 (*73*), following its recommended procedures with default settings. Recombination rates along the chromosomes were smoothed using the locally weighted scatterplot smoothing (LOWESS) method, utilizing the lowess function available in R version 4.3.3 (*74*) .

Characterizing TSDs and Protein Domains in LTR/Gypsy and LINE/L2

To investigate the insertion sites of LTR/Gypsy (Gypsy), we retrieved both the left and right target site duplications (TSDs) from the output of LTR_retriever. A custom Python script was then used to analyze sequences 30 base pairs upstream and downstream of the TSDs and calculated the nucleotide composition at each position.

To determine whether LINE/L2 (L2) contain TSDs of Gypsy, we followed the protocol of Goubert, Craig, Bilat, Peona, Vogan and Protasio (*75*) to retrieve full-length L2. In brief, we first blast the young L2 sequences (Kimura distance < 5%) obtained from the TEs annotation step, against the reference genome. We then aligned the L2 sequences using MAFFT (*76*), generated a consequence sequence using *cons* script from EMBOSS (*77*), and manually trimmed the TSDs using the visualization tool AliView (*69*). Sequentially, we used the script *getorf* to identify the potential open reading frames (ORFs), and then using InterProScan (*78*) to annotate the protein domains.

PIWI-RNA complex model prediction

To investigate the potential impact of mutations in the PAZ domain, we employed AlphaFold 3 (*79*) to build the model of protein structures and predict protein-RNA complex. The PIWI protein sequence was obtained from the previous gene prediction of Cb and identified through BLAST analysis against zebrafish PIWI1 (NP_899181.1, Ziwi). The domain annotation of the protein is based on UniProt(Q8UVX0). The RNA sequence was retrieved from the PDB database (pdb:3O7V). The protein structure was visualized by PyMOL (*80*) with very low confident region removed. The model prediction was highly confident and accurate (Fig. S20).

Migration rate estimation

We used gImble (*24*) to investigate the impact of chromosome fission on gene flow, particularly focusing on migration rates between Zp and Oe, as well as between Zp and Op. Two individuals were included for each species. Following the guidelines outlined in the gImble protocol, we used *bwa mem* to align reads from various samples to the reference genome of Zp. BAM files generated for the different comparisons were called for variants and subsequently merged into a single VCF file using the *freebayes-parallel* script. We prepared the dataset for gImble analysis with the *gimbleprep* script, and tested various models, including strict isolation (DIV), migration only (MIG), and isolation with migration (IM), using only the intergenic regions. Since gImble assumes a constant rate of unidirectional gene flow, the migration model can be further divided by species A to species B (IM_AB and MIG_AB) and species B to species A (IM_BA and MIG_BA).

Single copy gene and genetic distance

We used Orthofinder 2.5.4 (*81*) to identify single-copy genes (SCGs) between Zp and each of Cb, Op, Oe, and Ob. For Zp and Cb, we found 13,719 SCGs (Fission = 9,123, Nonfission = 4,596). Zp and Op yielded 13,604 SCGs (Fission = 8,401, Nonfission = 5,203), Zp and Oe resulted in 12,358 SCGs (Fission = 8,200, Nonfission = 4,158), and Zp and Ob had 13,880 SCGs (Fission = 9,220, Nonfission = 4,660). These SCGs were aligned using MACSE v2.06 (*82*). The p-distance was calculated using a custom Python script, which divided the number of nucleotide differences by the total number of nucleotides compared.

**Supplementary Text**

Genome assembly statistics of four Opsariichthyini species

The estimated genome size of Cb is 839.85Mb (Table S2-S3). The draft genome assembly by *Flye* was composed of 1,739 contigs and approximately 826.90Mb in genome size. The draft genome had an N50 of 2.70 Mb, and the largest contig is only 15.73Mb. The 1,739 contigs of polished draft genome were anchored into 1,233 contigs with a total length of 829.68Mb by Hi-C data. Among 1,233 contigs, the top 24 chromosome-level scaffolds were significantly larger than the rest contigs, which is consistent with the karyotype analyses of Cb (Fig. 1). These 24 pseudo-chromosomes ranged in size from 23.02 Mb to 46.66 Mb. N50 increased to 32.96Mb and L90 is 22. To validate the completeness of assembly, we mapped the short reads to the genome and got a 99.32% mapping rate, with an average coverage of 26x depths. Furthermore, the complete BUSCO score is 97.97% (Table S4).

The estimated genome size of Zp is 831Mb. The draft genome assembly was composed of 2,543 contigs and approximately 844.19Mb in genome size. The draft genome had an N50 of 6.17Mb, and the largest contig is only 18.26Mb. The 2,543 contigs of polished draft genome were anchored into 2,332 contigs with a total length of 847.09Mb by Hi-C data. We retrieved 24 pseudo-chromosomes, which consisted with literature. These 24 pseudo-chromosomes ranged in size from 23.38Mb to 46.34Mb and N50 was increased to 31.42. The mapping rate is 99.74%, with an average coverage of 34x depths. BUSCO score is 97.55%.

As for Oe, the draft genome assembly was composed of 2,931 contigs and approximately 919.07Mb in genome size. The draft genome had an N50 of 0.95Mb. The 2,931 contigs of polished draft genome were anchored into 2,835 contigs with a total length of 920.54Mb by Hi-C data, and therefore formed 39 pseudo-chromosomes. These pseudo-chromosomes ranged from 7.87Mb to 40.09Mb. N50 was increased to 21.75Mb and L90 is 38. The mapping rate is 99.31%, with an average coverage of 26x depths. The complete BUSCO score is 96.18%.

Finally, for Op, the draft genome assembly was composed of 2,749 contigs and approximately 915.44Mb in genome size. The draft genome had an N50 of 1.54Mb. The 2,749 contigs of polished draft genome were anchored into 2,483 contigs with a total length of 916.58Mb by Hi-C data, and thus formed 38 pseudo-chromosomes. These pseudo-chromosomes ranged from 8.07Mb to 42.83Mb. N50 was increased to 21.86Mb and L90 is 37. The mapping rate is 99.82%, with an average coverage of 15x depths. The complete BUSCO score is 97.69%.

Transposable elements (TEs) account for 51% to 55% of the opsariichthyin genomes, with DNA transposons being the most prevalent (~25 - 27%), followed by long terminal repeats (LTRs; ~7 - 8%), long interspersed nuclear element (LINEs; ~4 - 5%), and short interspersed nuclear element (SINEs; ~0.2 - 0.3%) (Table S5). Notably, the Opsariichthyini exhibit a significantly higher proportion of LINEs, particularly L2 elements (L2), and a significant increase in Gypsy elements (Gypsy) within their LTR content compared to other cyprinids. Gene annotation statistics reveal 26,894 to 30,730 protein-coding sequences in five opsariichthyin species, with BUSCO scores ranging from 91.10% to 96.70% (Table S6).

Centromere identification

To investigate the mechanism of chromosome fission, we identified putative centromere regions by searching for high order repeats (HORs) (*67*) and estimating recombination rates along the chromosomes. We discovered several HORs in Cb, Zp, and genus *Opsariichthys*. These sequences vary in length (Fig. S7A) but are composed of highly similar repeating units, each approximately 100 bp long with low GC content (Fig. S7A and S7B). Phylogenetic analysis suggests these HORs likely originated from an ancient LINE1 element (Fig. S7C). Notably, these HORs were predominantly found in regions of low recombination rates and gene density (Fig. S5-S6 and Fig. S8-S10), which also enriched with L2 and Gypsy. These features suggest that these regions may be putative centromeres, with HORs serving as centromeric DNA.

For Cb and Zp, the putative centromeres are characterized by regions containing centromeric DNA, low recombination rates, and enriched with L2 and Gypsy (Fig. S5 and S6). Based on the locations of these putative centromeres, we have categorized the chromosomes into metacentric, sub-metacentric, and sub-/telocentric categories (*15*). With the exception of one sub-metacentric chromosome in Cb and two sub-metacentric chromosomes in Zp, which were classified as sub-telocentric, the counts of the remaining chromosome types from the *in silico* analysis are consistent with those derived from karyotypic analysis, demonstrating the authenticity of our *in silico* karyotyping method based on the identification of putative centromeres.

We then searched for the centromere regions in *Opsariichthys* based on the following criteria: the presence of centromeric DNA, low recombination rate, high L2 and Gypsy, low gene density. Not all centromere regions defined in Cb and Zp met all these criteria comprehensively; therefore, we adopted a holistic approach to define the centromere regions in Opsariichthys, considering the overall genomic context. For Op, only three chromosomes derived from *in silico* and microscopic analyses were different (Table S1; Fig. S8). The karyotypes of Oe derived from *in silico* and microscopic analyses are identical (Fig. S9). In general, the results derived from different methods are largely consistent. We also conducted *in silico* karyotyping on the downloaded Ob genome (Fig. S10), which is identical with Oe.

Within *Opsariichthys*, approximately 83% ((28 + 33 + 35) / (39 + 38 + 39)) of chromosomes contain the centromere markers identified. In addition, in approximately 71% ((23 + 28 + 31) / (39 + 38 + 39)) of the chromosomes, the centromere markers are enriched in the putative centromere regions (Table S9-11). Furthermore, about 2/3 of fissioned chromosomes contain centromere markers in the putative centromere regions. This result support our hypothesis that fission events occurred within centromere regions, allowing each resultant chromosome to retain a partial centromere necessary for proper segregation.

While centromere markers are present in almost all centromeres of Cb and Zp, only about 70% of putative centromeres in *Opsariichthys* species show these markers. This discrepancy likely arises because *Opsariichthys* predominantly features sub-/telocentric chromosomes. Assembling telomeric regions is challenging due to their repetitive sequences, which can cause sequencing errors and hinder accurate assembly. In comparison, 97% of contig lengths were anchored to pseudo-chromosomes in Cb, whereas only 91% reached this stage in Op and Oe.

Comparative Analysis of Piwi1 Protein Across Species

To investigate the Piwi1 protein across species, we performed a BLAST search using the zebrafish Piwi1 protein (NP_899181.1) as a query. This analysis was conducted against the protein-coding genes of various species, utilizing the annotated protein domain structure of NP_899181.1 as a reference. In the tribe Opsariichthyini, the Piwi1 protein comprises 859 amino acids, identical to that of zebrafish. Compared to NP_899181.1, the Piwi1 protein in Opsariichthyini species exhibits 36 to 40 mutations. Within the PAZ domain (positions 275–412), 7 to 9 amino acid differences were observed. However, only two mutations were shared by all Opsarichthyini species, G348S and V380I. All PAZ domain sequences used in this study can be found at the end of *Supplementary Texts*.

TEs in the breaking region Macaque chromosome 7

The presence of active TEs associated with chromosome fission has also been observed in primate evolution. Human chromosomes 14 and 15 resulted from a single fission event of a hominoid ancestral chromosome, which most closely resembles macaque chromosome 7 (*83*). Detailed analysis of the fission site identified LINE1 (L1) as being associated with chromosome breakage (*84*). We found that the breaking region of macaque chromosome 7 spans approximately 300 kb (Fig. S21A). Within this region, the density of L1 is nearly three times higher compared to the rest of the chromosome (Fig. S21B). Additionally, this region has experienced a recent L1 expansion (Fig. S21C).

Frequent fission and fusion in *Opsariichthys* genomes

The evidence suggests that chromosome fission and fusion occurred frequently among species of *Opsariichthys*, providing numerous opportunities for accumulation of multiple fissions within their populations, thereby enhancing the potential for speciation through increased genetic diversity across different localities. For example, a recent study revealed that the karyotypes of Ob from Yangtze River, Pearl River, and Qiantang River basins were N = 37, 38, and 39, respectively (*85*). In addition, chromosome 16 of Zp was shown to split into two separate chromosomes in Oe and Ob (Fig. 2; Table S13), whereas its homolog in Op, chromosome 9, did not undergo fission. Given the close relationship between Op and Ob (Fig. 2), it is likely that the fissions in Oe and Ob occurred independently after their divergence or that previous fissions fused again in Op, resulting in N =38. Either scenario implies frequent chromosome fission and fusion in *Opsariichthys*.

All PAZ domain of piwi1 protein for all species in this study

>Drer_PAZ

ETVLDFMYSLRQQCGDQRFPEACTKELVGLIILTKYNNKTYRIDDIAWDHTPNNTFKKGDTEISFKNYFKSQYGLDITDGNQVLLVSHVKRLGPSGRPPPGPAMLVPEFCYLTGLTDKMRADFNIMKDLASHTRLSPE

>Dtra_PAZ

ETVLDFMGTLKQQCGDQRFIEACTKELVGLIILTKYNNKTYRIDDIAWDHNPSNTFKKGDAEISFKDYFKTQYGLNVTDGYQALLVSRVKRLGPSGQPPPGPALLVPEFCYLTGLTDKMRADFNIMKDLATHTRLSPE

>Lroh_PAZ

ETVLDFMYSLRQQ----------------------YNNKTYRIDDIAWDHTPNNTFKKGDTEISFKNYFKTQYGLDITDGNQVLLVSHVKRLGPS??-----------------LTDKMRADFNIMKDLASHTRLSPE

>Sgra_PAZ

ETVLDFMYSLRQQCGDQRFPEACTKELVGLIILTKYNNKTYRIDDIAWDHTPNNTFKKGDTEISFKNYFKTQYGLDITDGNQALLVSHVKRLGPSGRPPPGPAMLVPEFCYLTGLTDKMRADFNIMKDLASHTRLSPE

>Omac_PAZ

ETVLDFMYSLRQQCGDQRFPEACTKELVGLIILTKYNNKTYRIDDIAWDHTPNNTFKKGDTEISFKNYFKTQYGLDITDGNQVLLVSHVKRLGPSGRPPPGPAMLVPEFCYLTGLTDKMRADFNIMKDLASHTRLSPE

>Ptet_PAZ

ETVLDFMYSLRQQCGDQRFPEACTKELVGLIILTKYNNKTYRIDDIAWDHTPNNTFKKGDTEISFKHYFKTQYGLDITDGNQVLLVSHVKRLGPSGRPPPGPAMLVPEFCYLTGLTDKMRADFNIMKDLASHTRLSPE

>Ppro_PAZ

ETVLDFMYSLRQQCGDQRFAEACTKELVGLIILTKYNNKTYRIDDIAWDHTPNNTFTKGDTEISFKNYFKTQYGLDITDGNQALLVSHVKRPGPAGRPPPGPAMLVPEFCYLTGLTDKMRADFNIMKDLASHTRLSPE

>Agar_PAZ

ETVLDFMYSLRQQCGDQRFSEACTKELVGLIILTKYNNKTYRIDDIAWDHTPNNTFKKGDTEISFKNYFKTQYGLDITDGNQALLVSHVKRLGPSGRPPPGPAMLVPEFCYLT-----MRADFNIMKDLASHTRLSPE

>Mamb_PAZ

ETVLDFMYSLRQQCGDQRFSEACTKELVGLIILTKYNNKTYRIDDIAWDHTPNNTFKKGDTEISFKNYFKTQYGLDITDGNQALLVSHVKRLGPAGRPPPGPAMLVPEFCYLTGLTDKMRADFNIMKDLASHTRLSPE

>Cbar_PAZ

ETVLDFMYSLRQQCGEQRFSEACTKELVGLIILTKYNNKTYRIDDIAWDHTPNNTFKKGDTEISFKNYFKTQYSLDITDGNQALLVSHVKRLGPAGRPPPGPAMLIPEFCYLTGLTDKMRADFNIMKDLSLHTRLSPE

>Zpla_PAZ

ETVLDFMYSLRQQCGDHRFSEACTKELVGLIILTKYNNKTYRIDDIAWDHTPNNTFKKGDTEISFKNYFKTQYSLDITDGNQALLVSHVKRLGPAGRPPPGPAMLIPEFCYLTGLTDKMRADFNIMKDLASHTRLSPE

>Oevo_PAZ

ETVLDFMYSLRQQCGDHRFSEACTKELVGLIILTKYNNKTYRIDDIAWDHTPNNTFKKGDTEISFKNYFKTQYSLDITDGNQALLVSHVKRLGPAGRPPPGPAMLIPEFCYLTGLTDKMRADFNIMKDLASHTRLSPE

>Obid_PAZ

ETVLDFMYSLRQQCGDHRFSEACTKELVGLIILTKYNNKTYRIDDIAWDHTPNNTFKKGDTEISFKNYFKTQYSLDITDGNQALLVSHVKRLGPAGRPPPGPAMLIPEFCYLTGLTDKMRADFNIMKDLASHTRLSPE

>Opac_PAZ

ETVLDFMYSLRQQCGDHRFSEACTKELVGLIILTKYNNKTYRIDDIAWDHTPNNTFKKGDTEISFKNYFKTQYSLDITDGNQALLVSHVKRLGPAGRPPPGPAMLIPEFCYLTGLTDKMRADFNIMKDLASHTRLSPE

>Ttib_PAZ

ETVLDFMYSLRQQCGDQRFPEACTKELVGLIVLTKYNNKTYRIDDIAWDQNPNNTFKRGDTDISFRNYFKTQYGLDITDGNQVLLVSHVKRLGPAGRPPPGPALLIPEFCYLTGLTDKMRSDFNIMKDLATHTRLSPE

>Tble_PAZ

ETVLDFMYSLRQQCGDQRFPEACTKELVGLIVLTKYNNKTYRIDDIAWDQNPNNTFKRGDTDISFRNYFKTQYGLDITDGNQVLLVSHVKRLGPSGRPPPGPALLIPEFCYLTGLTDKMRSDFNIMKDLATHTRLSPE

**Supplementary Figures**

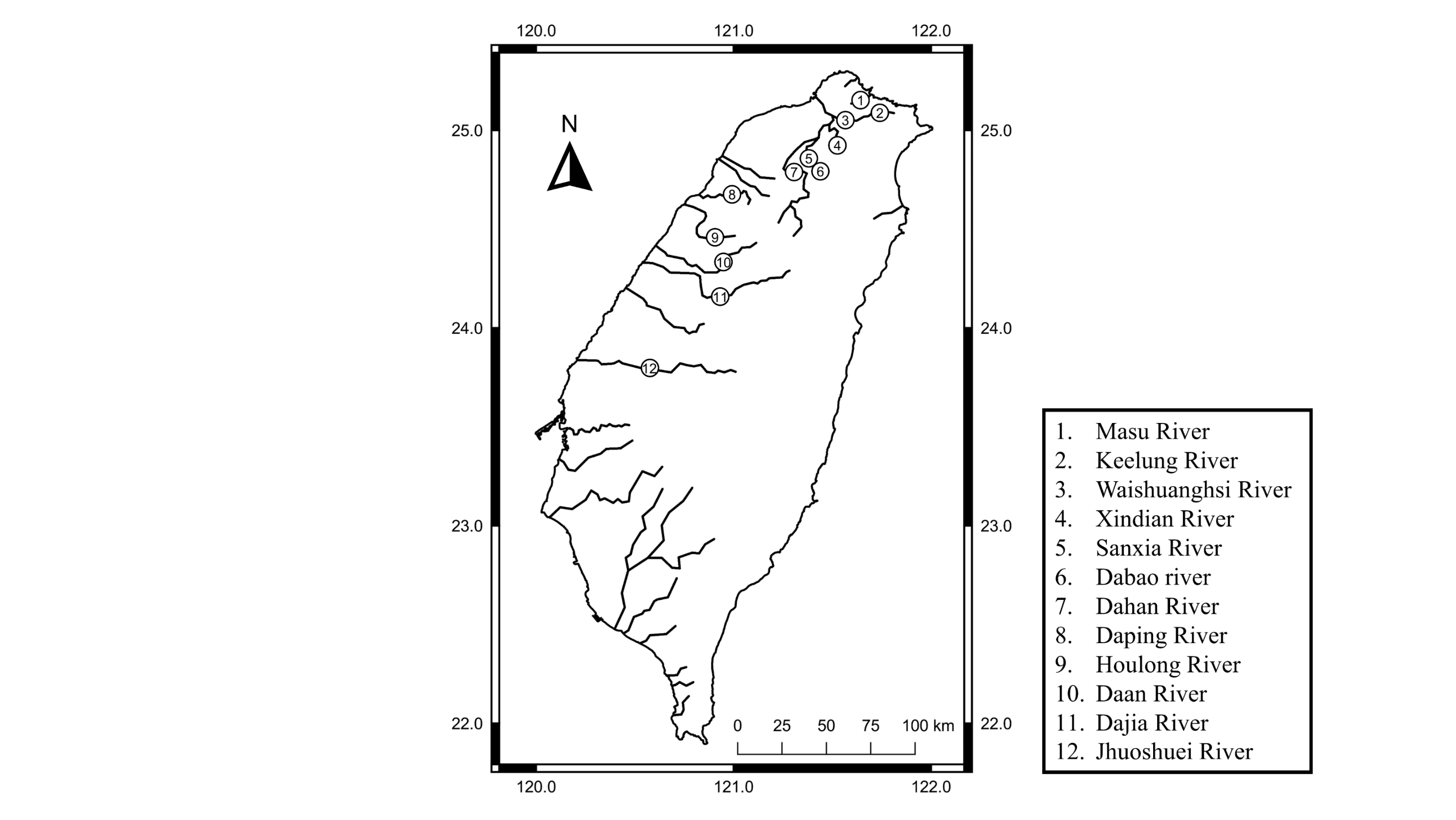

**Fig. S1. Sampling locations map.**

The map shows the sampling locations in Taiwan. The rivers sampled are numbered from north to south as follows: 1) Masu River, 2) Keelung River, 3) Waishuanhsi River, 4) Xindian River, 5) Sanxia River, 6) Dabao River, 7) Dahan River, 8) Daping River, 9) Houlong River, 10) Daan River, 11) Dajia River, 12) Wu River, 13) Jhoushuei River and 14) Puzih River (See also Supplementary Table 9).

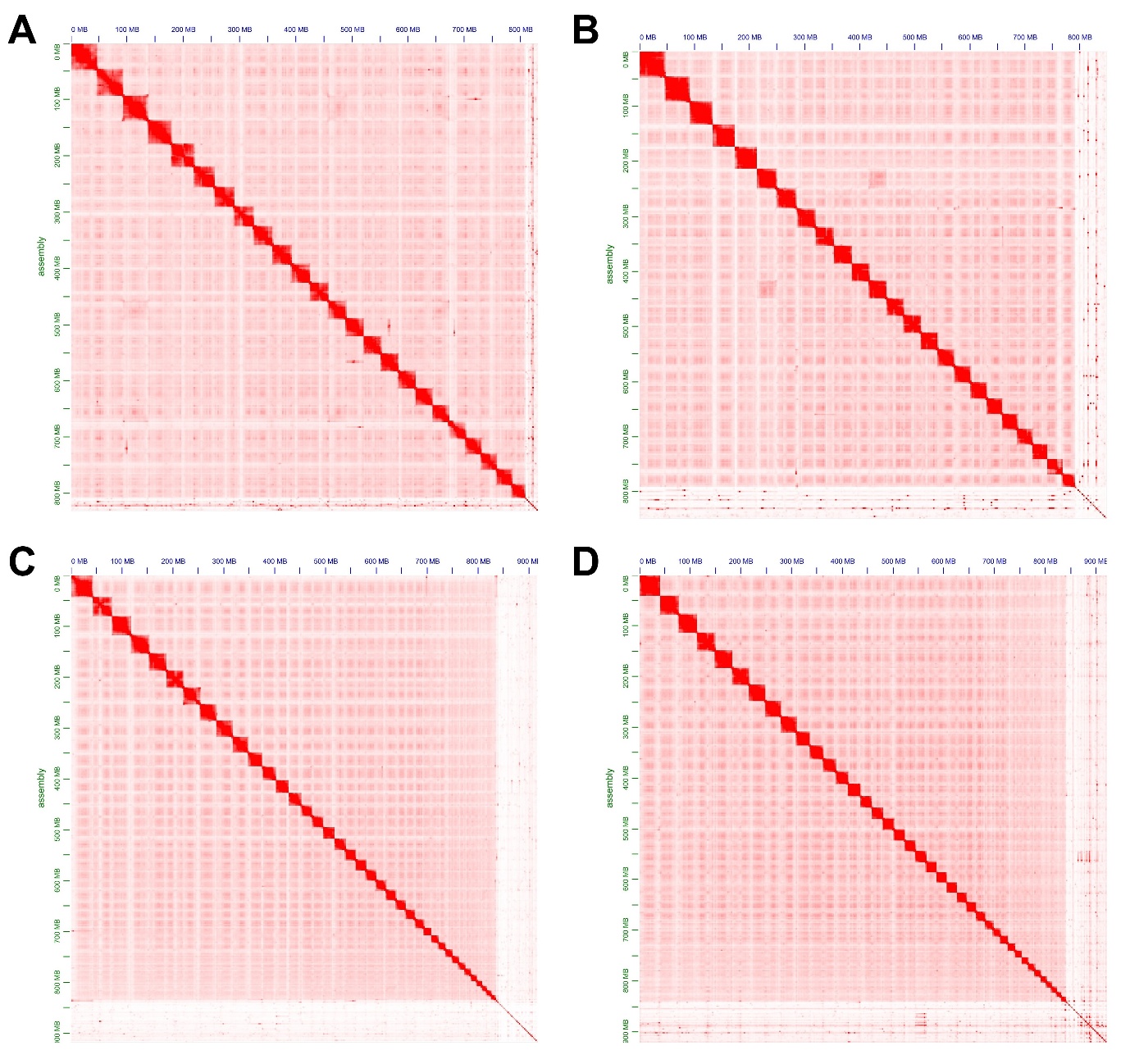

**Fig. S2. Heat map of interactive intensity between chromosome sequences of A,** Cb (*Candidia barbatus*) **B,** Zp (*Zacco platypus*) **C,** Op (*Opsariichthys pachycephalus*), and **D,** Oe (*O. evolans*) anchored by Hi-C.

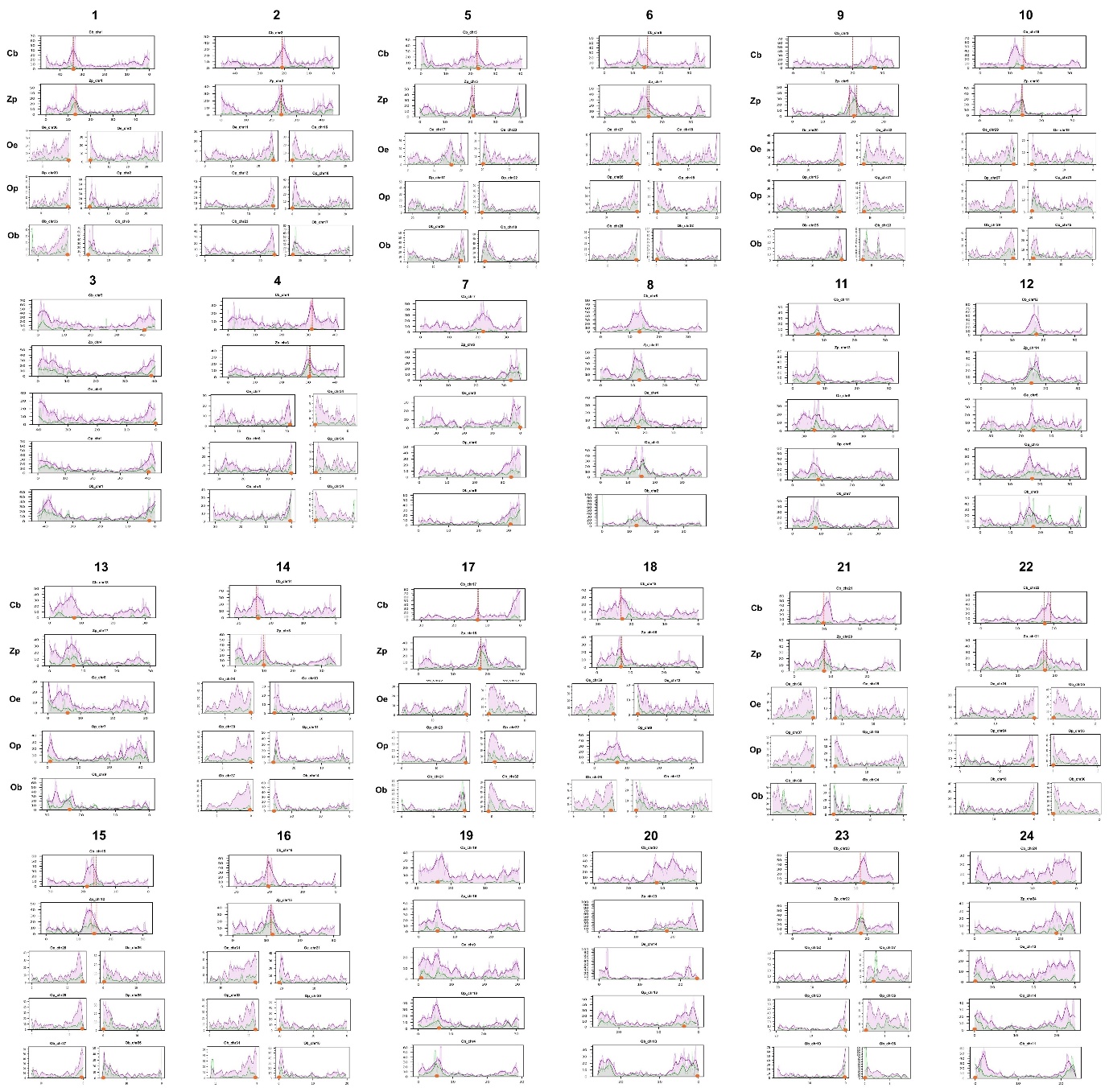

**Fig. S3. Syntenic relationships across studied species for each chromosome.**

The dashed line in Cb (*Candidia barbatus*) and Zp (*Zacco platypus*) indicated the breaking regions. The orange solid dot represents the putative centromere region. The young L2 and Gypsy (k-distance < 5%), shown in purple and green respectively, are predominantly localized around putative centromeres and breaking regions.

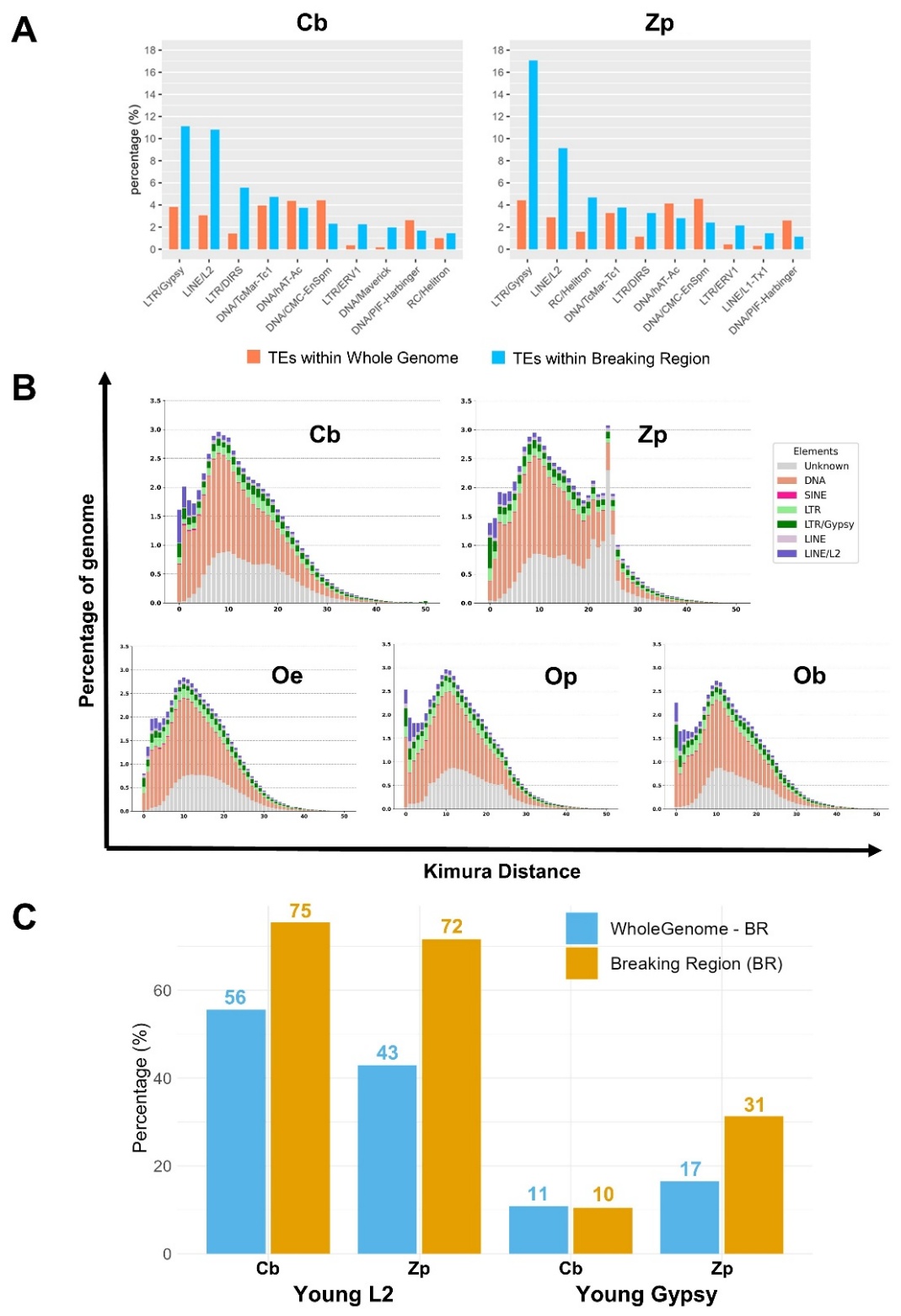

**Fig. S4. Distribution of transposable elements in the opsariichthyini** **genomes.**

**A,** Proportion of different TEs in the whole genomes and breaking regions. In Cb (*C. barbatus*), the proportions of L2 and Gypsy in the breaking regions are 3.5- and 2.9-fold higher, respectively, than in the entire genome. In Zp (*Z. platypus*), they are 3.2- and 3.9-fold higher, respectively. **B,** Repeat landscapes among five opsariichthyini species. The y-axis represents the percentage of the genome occupied by each repeat element, and the x-axis represents the sequence divergence (measured by Kimura distance). **C,** Young L2 (Kimura distance < 5%) is enriched within the breaking regions of both Cb and Zp. In contrast, recent expansion of Gypsy within the breaking regions is more pronounced in Zp.

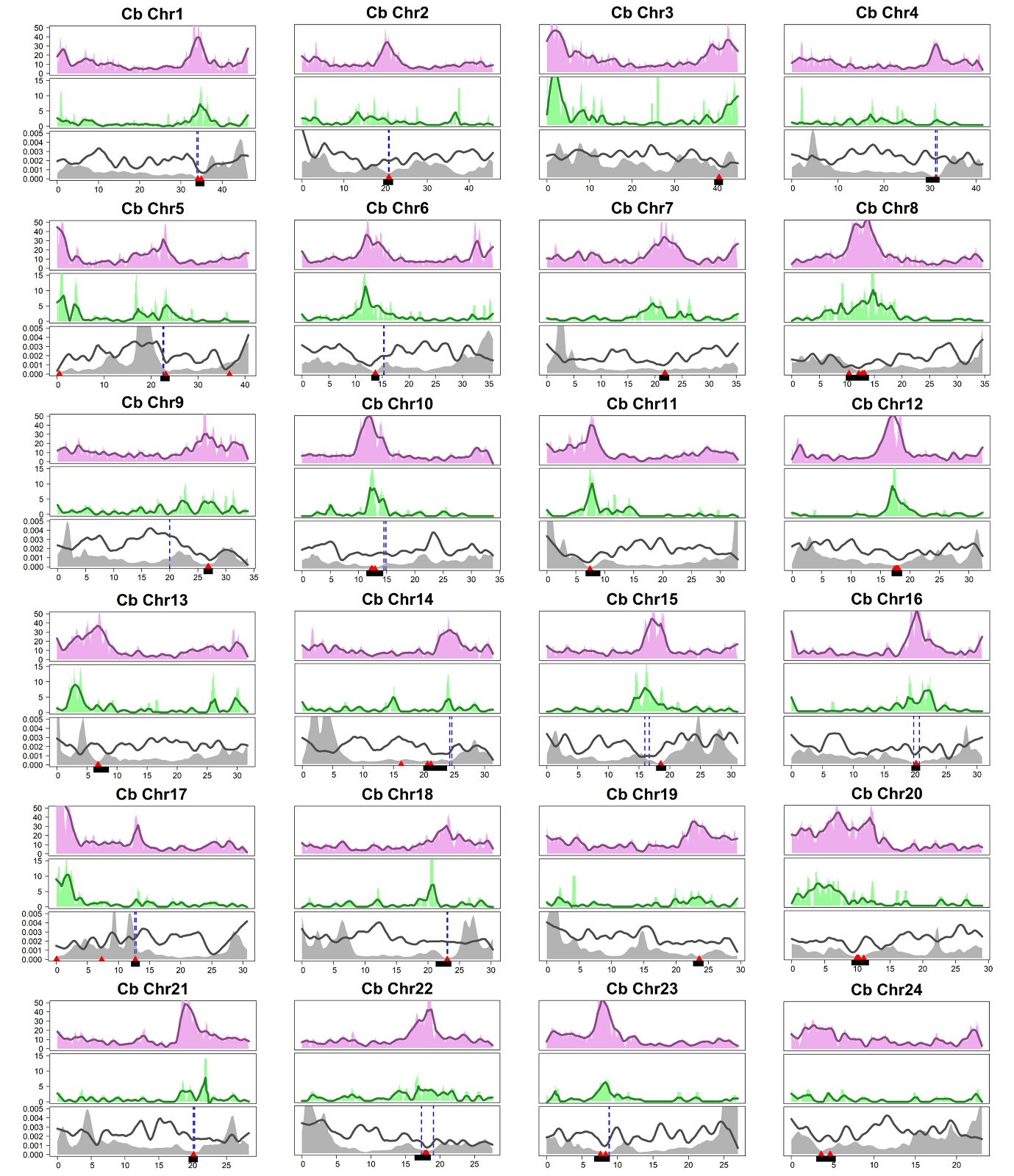

**Fig. S5. Distribution of L2, Gypsy, gene density, and recombination rate on each chromosome of Cb (*Candidia barbatus*).**

**Upper panels:** Distribution of recent L2 (divergence < 5%). **Middle panels:** Distribution of recent Gypsy (divergence < 5%). **Lower panels:** Present recombination rates (shaded areas) and gene densities (black lines). Dotted blue lines indicate breaking regions. Red triangles represent centromere markers, and black line segments denote putative centromeres.

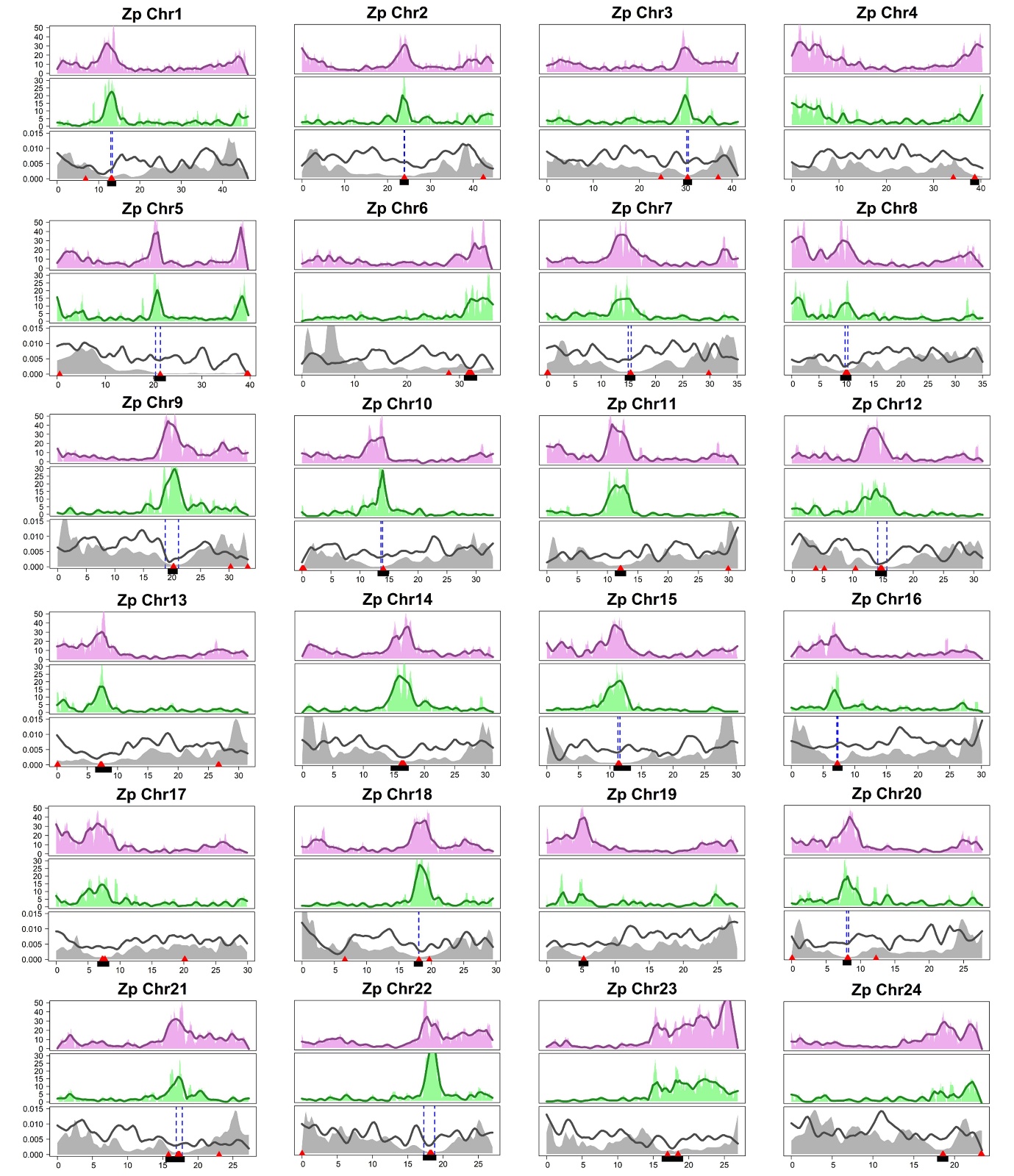

**Fig. S6. Distribution of L2, Gypsy, gene density, and recombination rate on each chromosome of Zp (*Zacco platypus*).**

**Upper panels:** Distribution of recent L2 (divergence < 5%). **Middle panels:** Distribution of recent Gypsy (divergence < 5%). **Lower panels:** Present recombination rates (shaded areas) and gene densities (black lines). Dotted blue lines indicate breaking regions. Red triangles represent centromere markers, and black line segments denote putative centromeres.

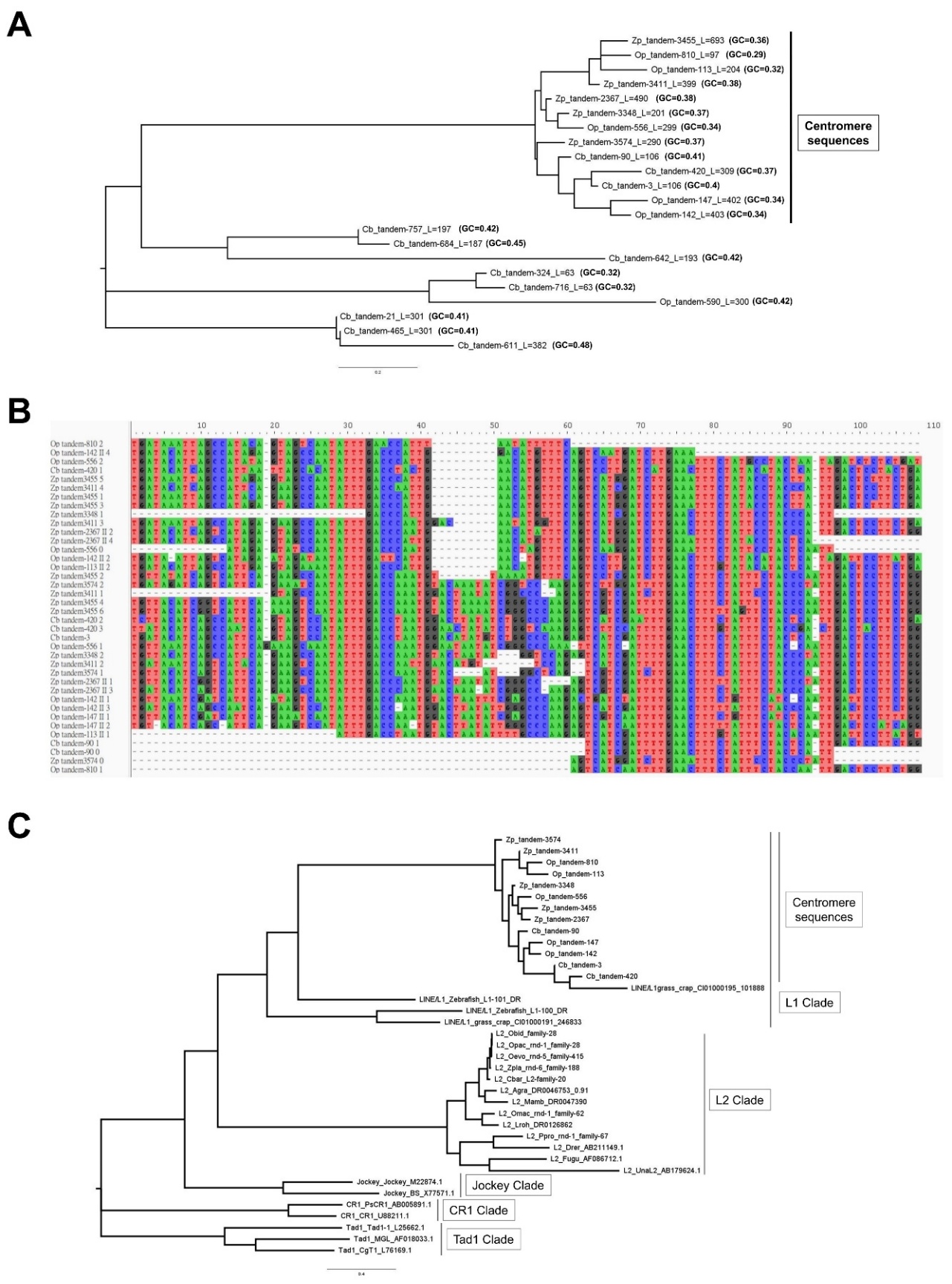

**Fig. S7. Centromere sequences of the Opsariichthyini.**

**A,** Phylogeny of high order repeats. Sequence name is present by species name, repeat ID, sequence length(L), and GC content. Lower GC content cluster is chosen as putative centromere sequences. **B,** Alignment of putative centromere sequences. Fragment of each centromere sequences were split manually, with a length approximately of 100 basepairs. **C,** Phylogeny of LINE elements that are related to centromere sequences.

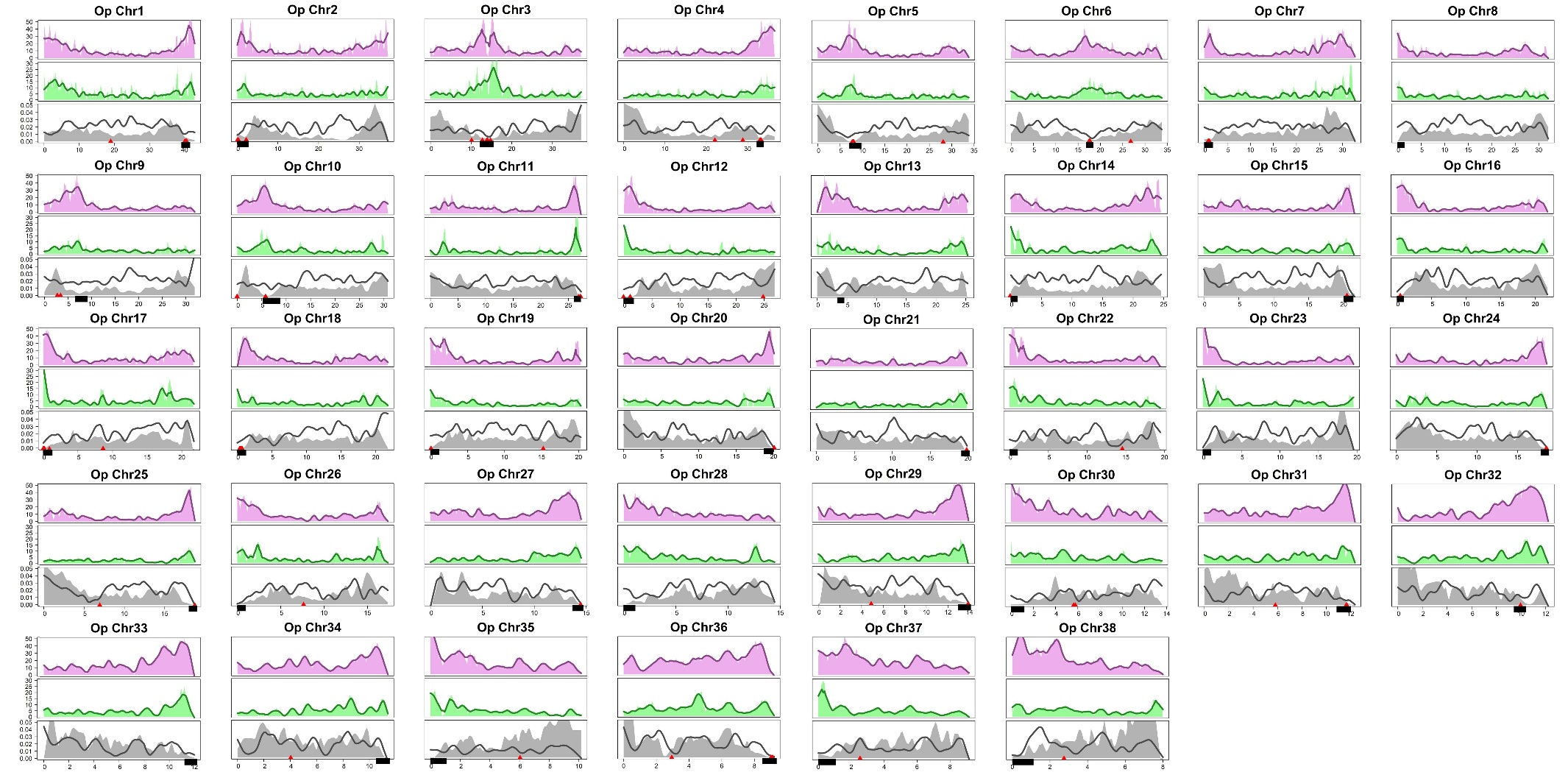

**Fig. S8. Distribution of L2, Gypsy, gene density, and recombination rate across chromosomes of Op (*Opsariichthys pachycephalus*).**

Upper panels: Distribution of recent L2 (divergence < 5%). Middle panels: Distribution of recent Gypsy (divergence < 5%). Lower panels: Present recombination rates (shaded areas) and gene densities (black lines). Dotted blue lines indicate breaking regions. Red triangles represent centromere markers, and black line segments denote putative centromeres.

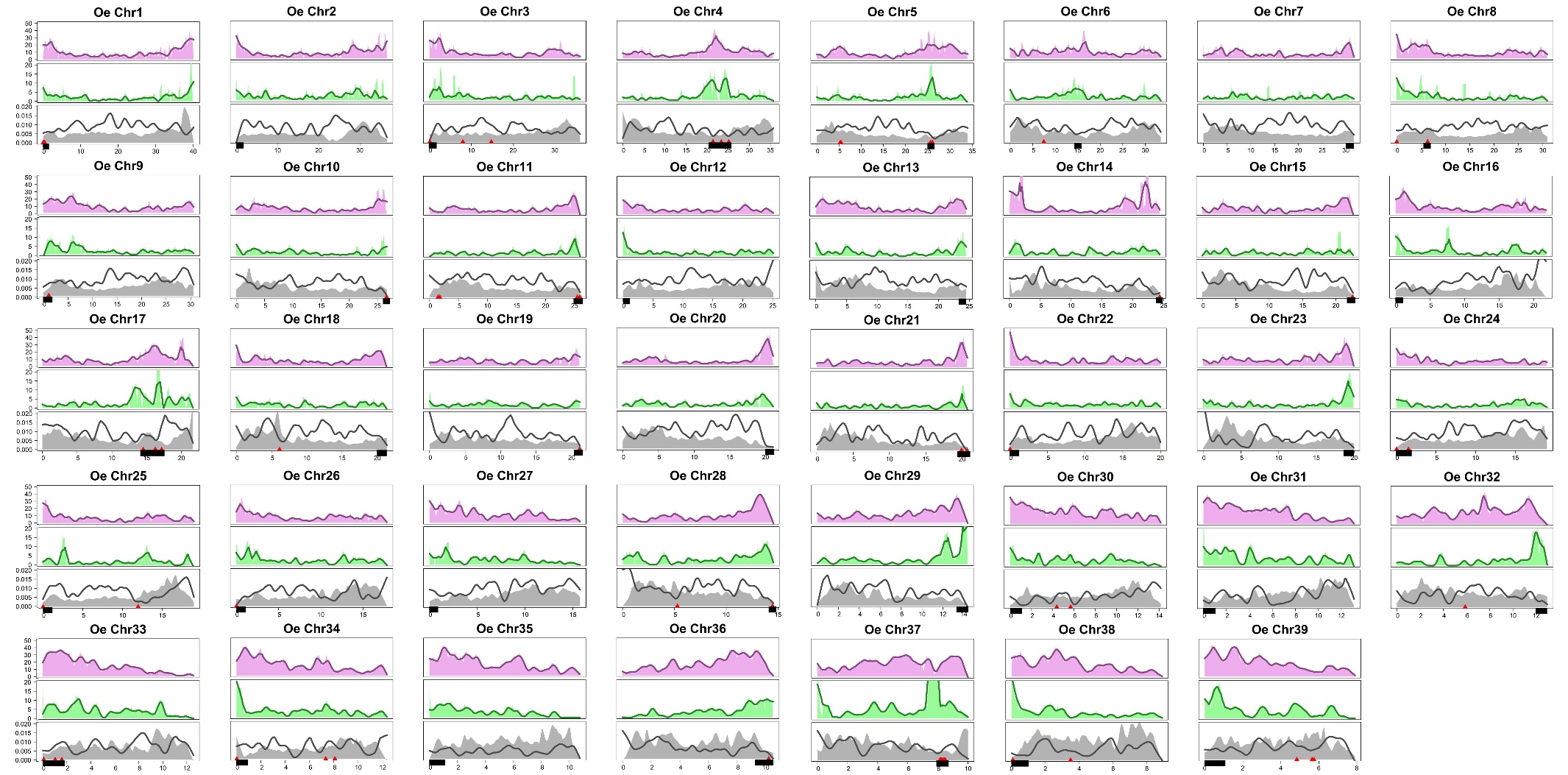

**Fig. S9. Distribution of L2, Gypsy, gene density, and recombination rate across chromosomes of Oe (*Opsariichthys evolans*).**

Upper panels: Distribution of recent L2 (divergence < 5%). Middle panels: Distribution of recent Gypsy (divergence < 5%). Lower panels: Present recombination rates (shaded areas) and gene densities (black lines). Dotted blue lines indicate breaking regions. Red triangles represent centromere markers, and black line segments denote putative centromeres.

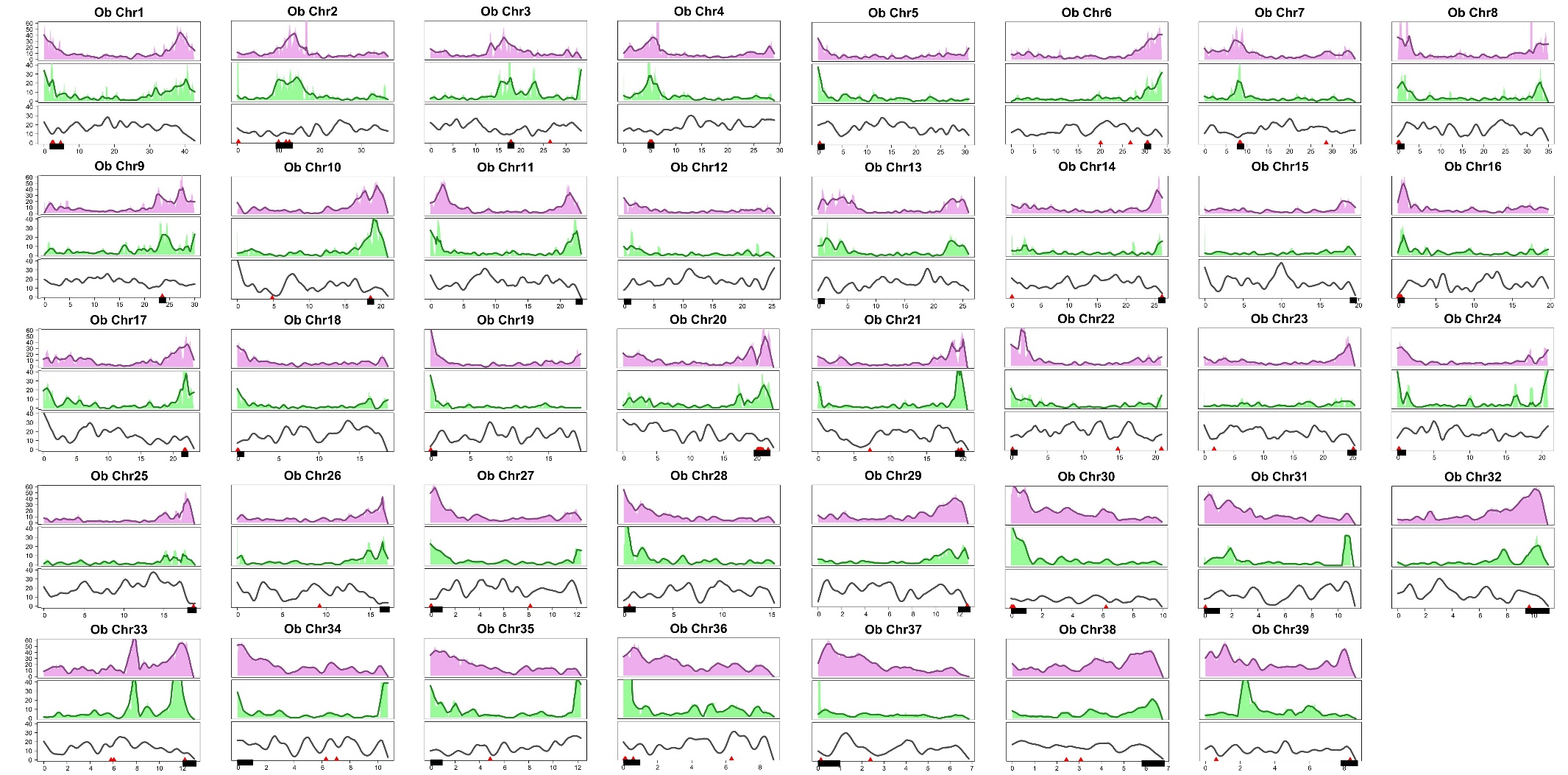

**Fig. S10. Distribution of L2, Gypsy, gene density, and recombination rate across chromosomes of Ob (*Opsariichthys bidens*).**

Upper panels: Distribution of recent L2 (divergence < 5%). Middle panels: Distribution of recent Gypsy (divergence < 5%). Lower panels: Present gene densities (black lines). Dotted blue lines indicate breaking regions. Red triangles represent centromere markers, and black line segments denote putative centromeres.

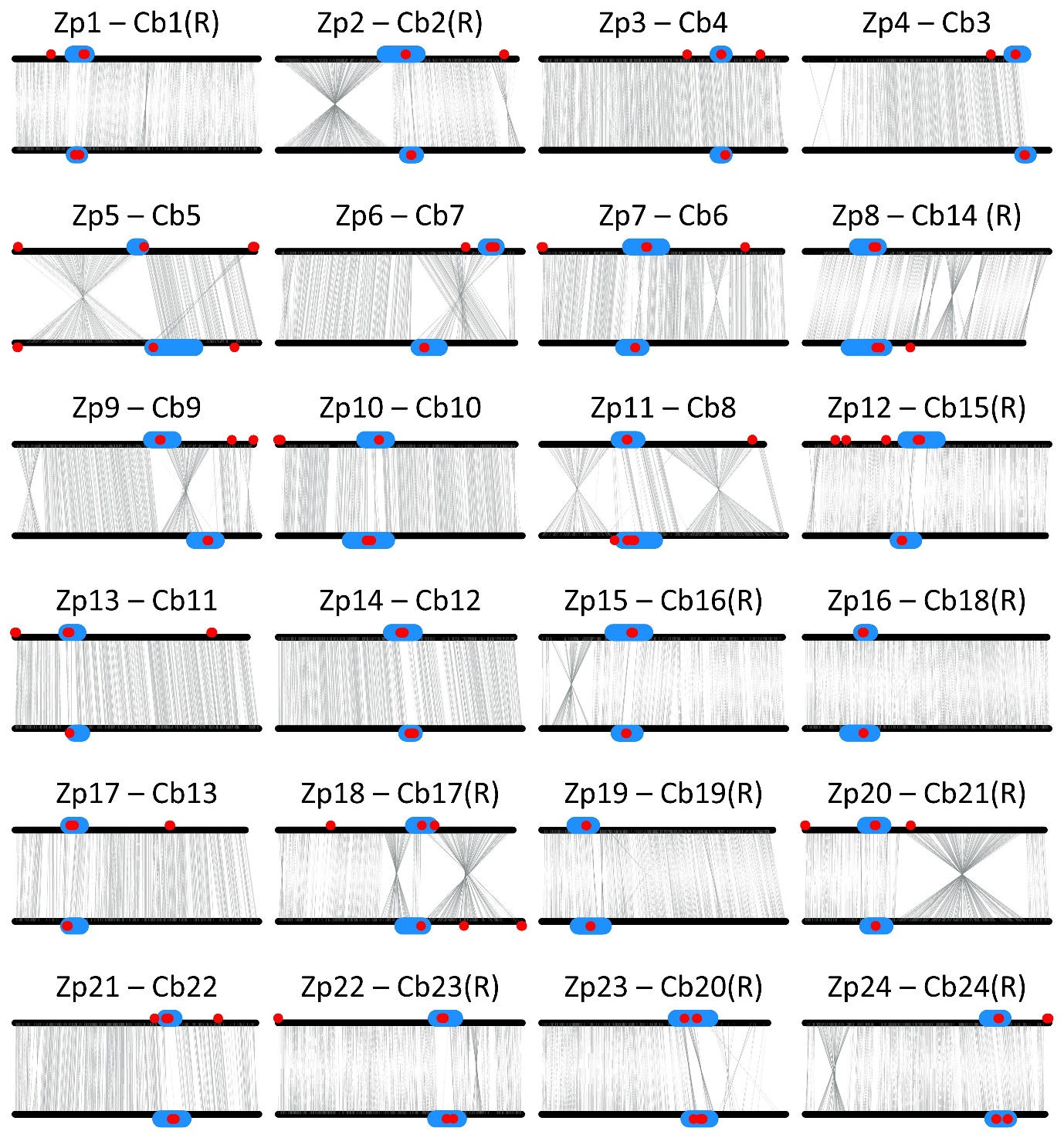

**Fig. S11. Syntenic relationships between Zp (above) and Cb (below) chromosomes highlight frequent pericentric inversions between species**

The red dots are the centromere markers, and blue regions represent the putative centromere regions. The orientations of synteny blocks are based on those in Zp. In Cb, the notation “(R)” indicates that the synteny orientation is reversed.

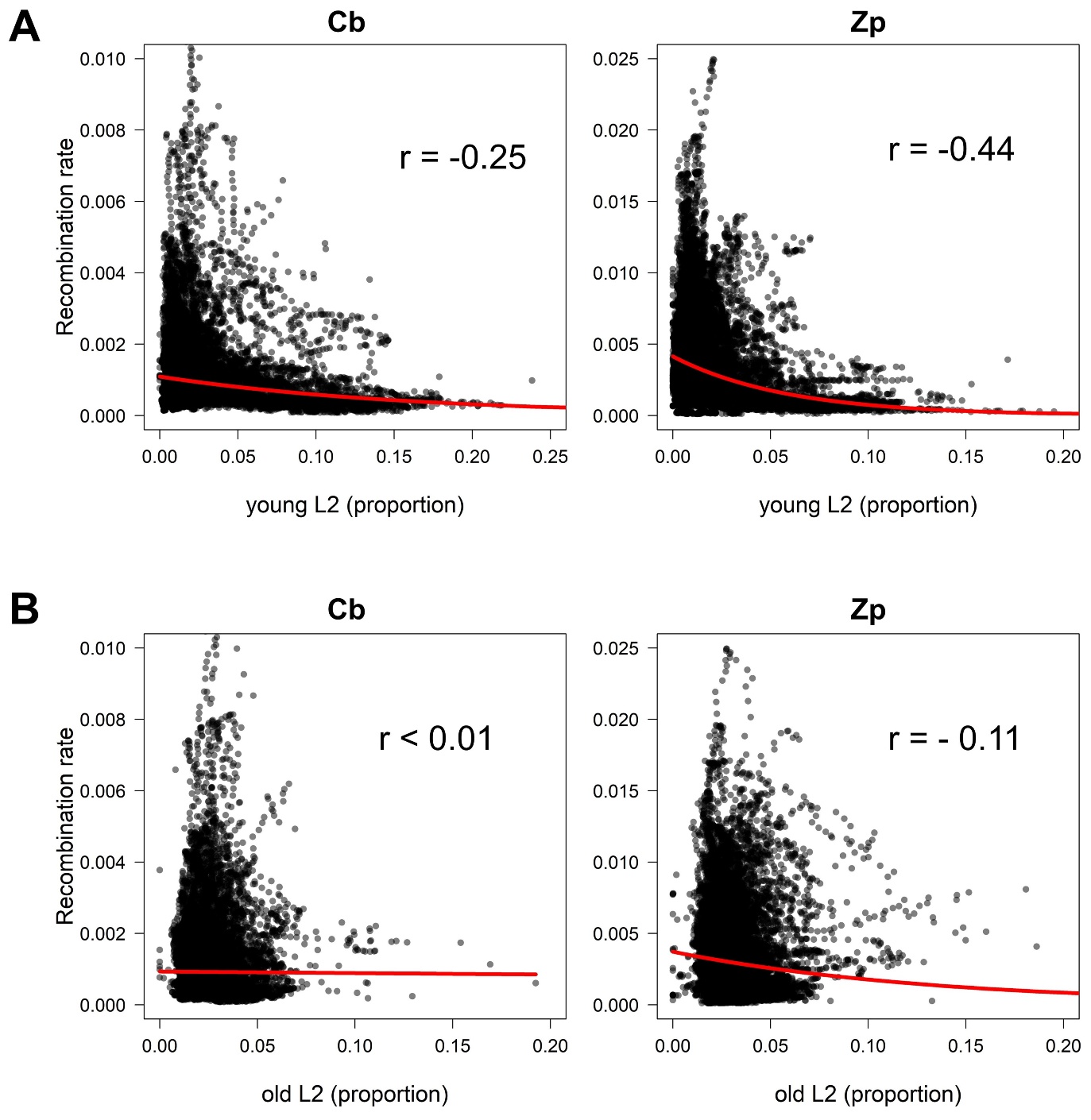

**Fig. S12. Correlation between L2 and recombination rate in Cb (*Candidia barbatus*, left panel) and Zp (*Zacco platypus*, right panel).**

**A,** Young L2 (K-distance < 5%) and **B,** Old L2 (K-distance > 5%)

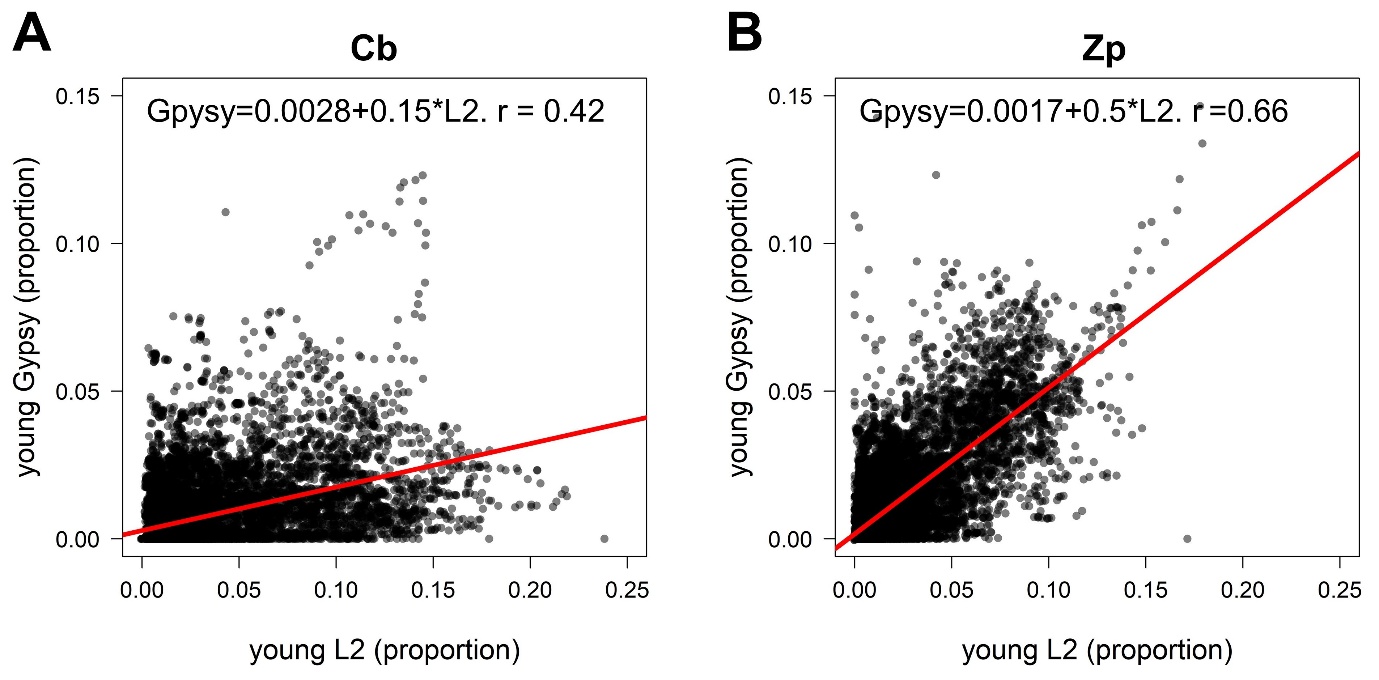

**Fig. S13. Correlation between young L2 and Gypsy in A, Cb (*Candidia barbatus*) and B, Zp (*Zacco platypus*).**

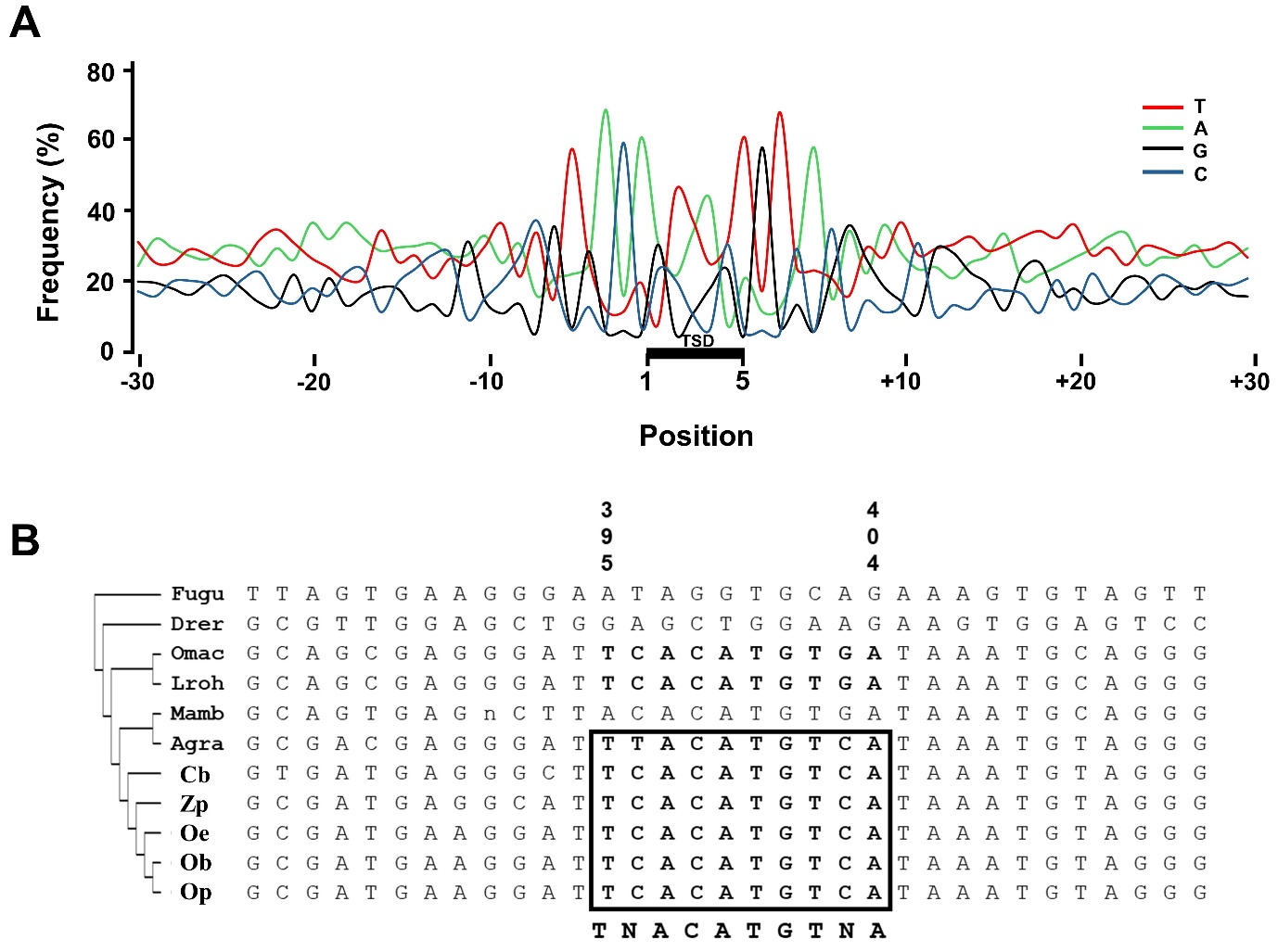

**Fig. S14. Insertion preference of Gypsy.**

**A,** Positions 1 to 5 represent the target site duplication (TSD) sequence. Negative and positive numbers indicate the distance in base pairs from the TSD. Frequencies of T(red), A(green), G(black) and C(blue) at each position are plotted. A conserved palindrome sequence flanking the 5-bp TSD in opsariichthyin species reads as TNACA-TSD-TGTNA. **B,** The alignment of L2 from different species. The insertion sites of Gypsy can be found in L2 of the Opsariichthyini (in box).

Fugu: *Takifugu rubripes*; Drer: *Danio rerio*; Omac: *Onychostoma macrolepis*; Lroh: *Labeo rohita*; Mmab: *Megalobrama amblycephala*; Agra: *Anabarilius graham*; Cb: *Candidia barbatus*; Zp: *Zacco platypus;* Oe: *Opsariichthys evolans;* Op: *O. pachycephalus;* Ob: *O. bidens*

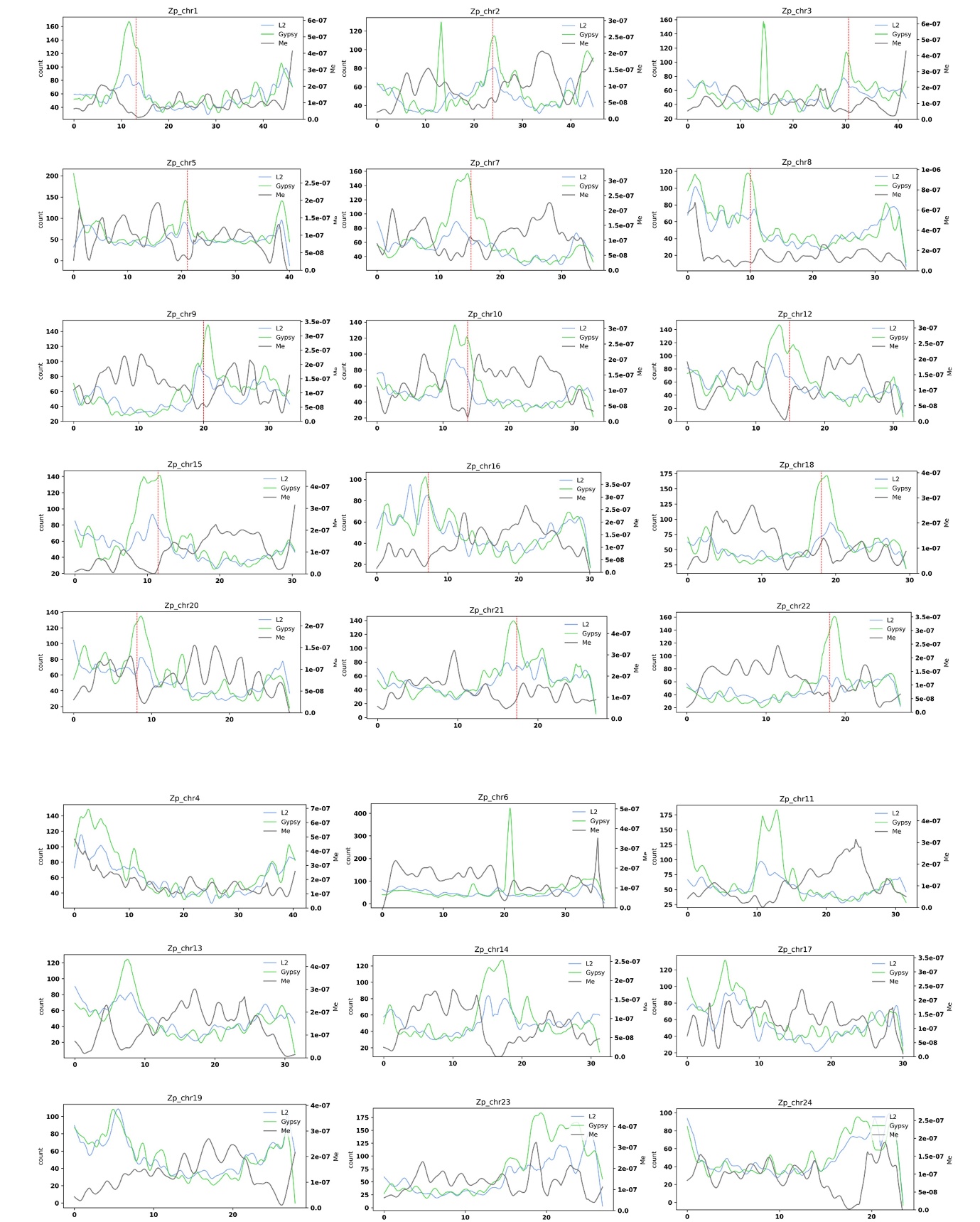

**Fig. S15. Migration rate (Me) between Zp (*Z. platypus*) and Oe (*O. evolans*) across various chromosomes.**

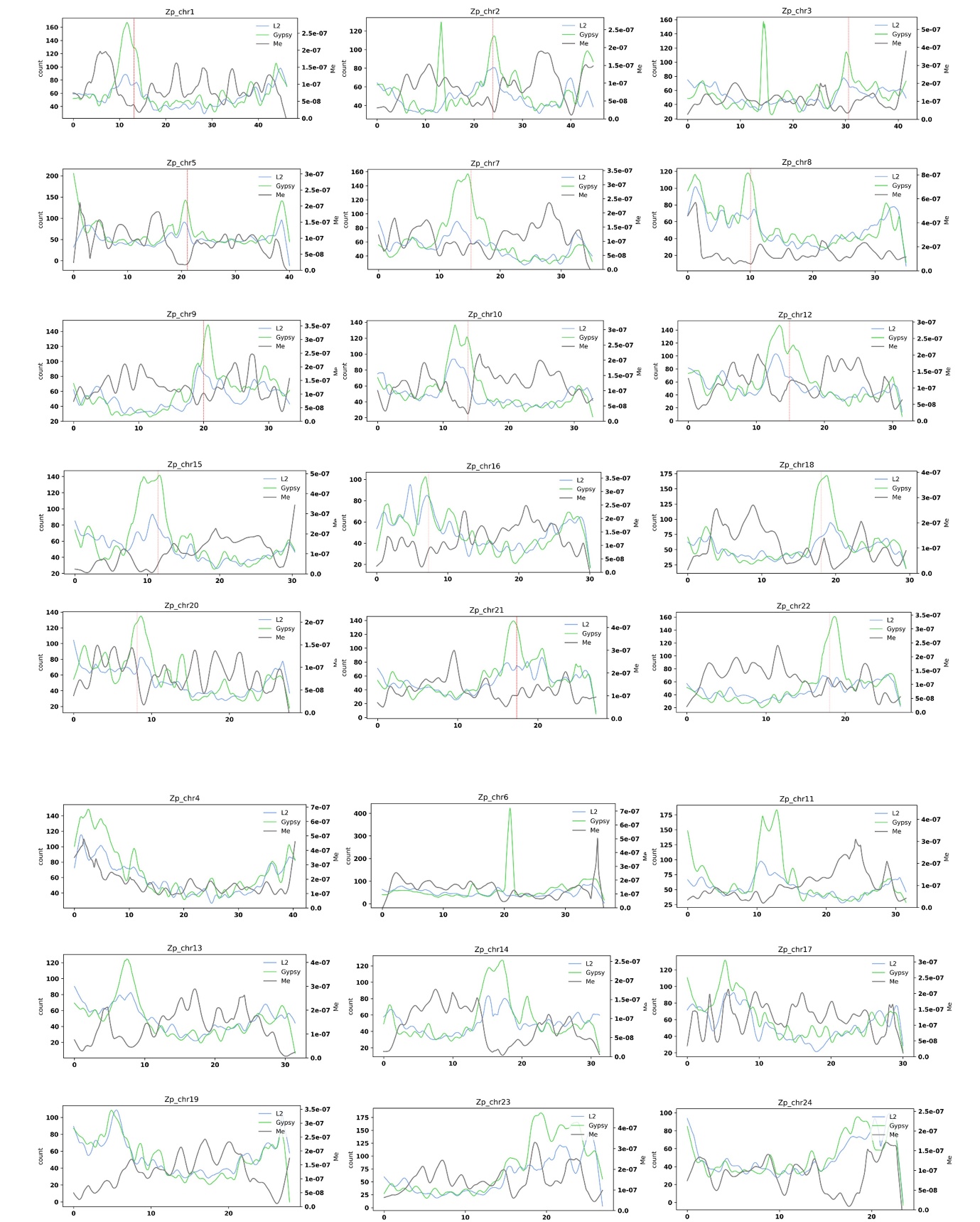

**Fig. S16.** **Migration rate (Me) between Zp (*Z. platypus*) and Op (*O. pachycephalus*) across various chromosomes.**

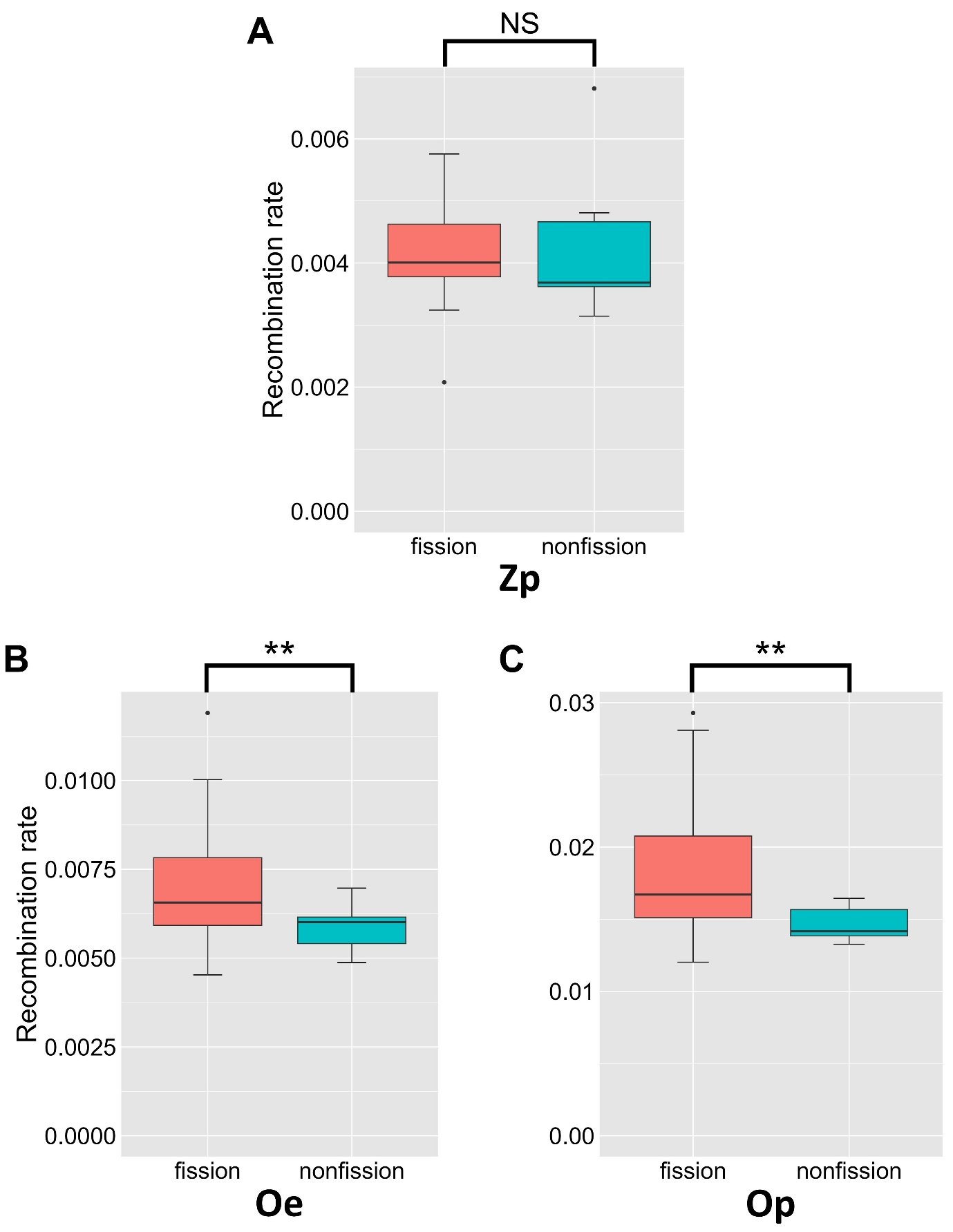

**Fig. S17.** **Recombination rate between fission and non-fission chromosomes.**

**A,** Zp (*Zacco platypus*) **B,** Oe (*Opsariichthys evolans*) and **C,** Op (*Opsariichthys pachycephalus*).

** p < 0.01

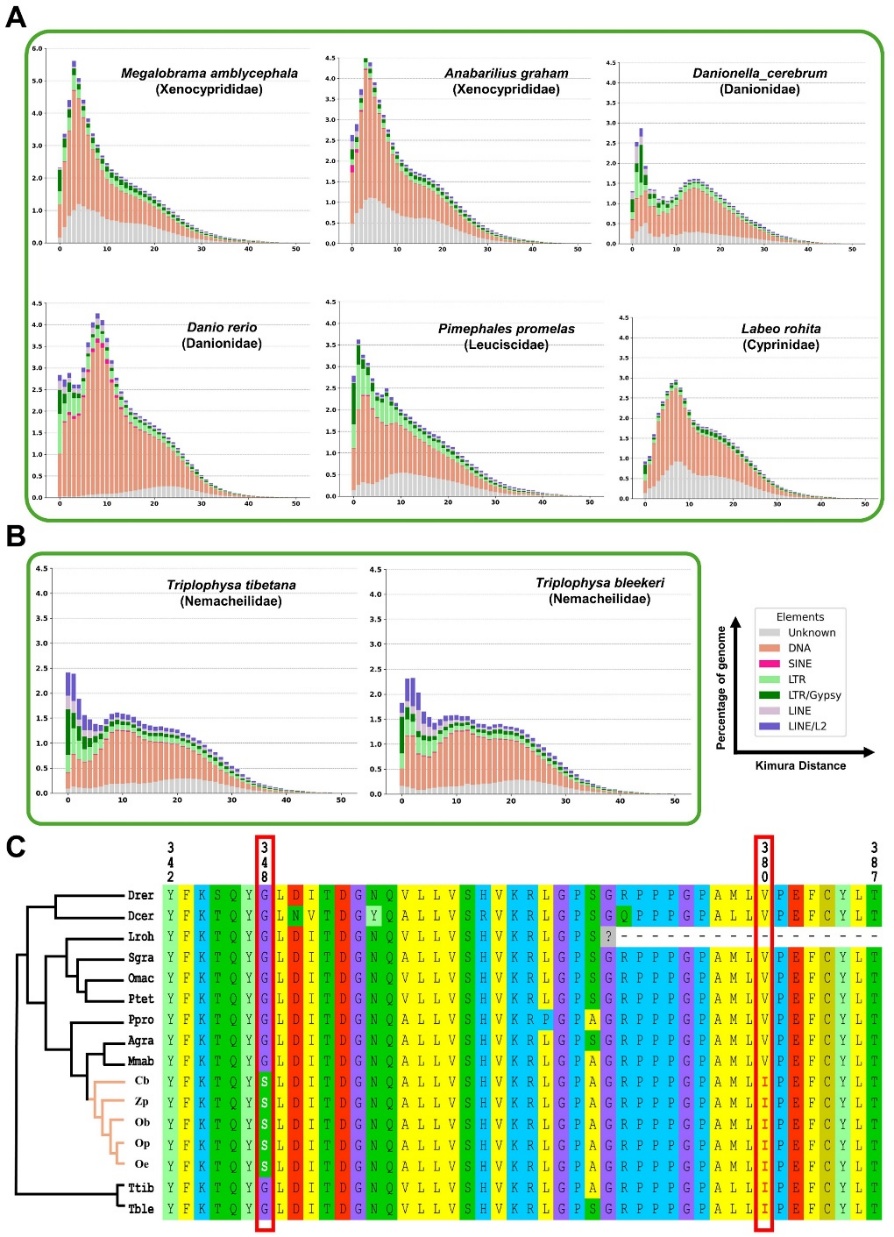

**Fig. S18.** **Association of mutations in PAZ domain of the PIWI1 protein with L2 expansion.**

Repeat landscapes of **A,** six Cyprinoidei species and **B,** two stone loach species. The y-axis represents the percentage of the genome occupied by each repeat element, and the x-axis represents the Kimura distance. DNA transposons are shown in orange, SINEs in red, LTRs (excluding Gypsy) in light green, Gypsy in green, LINEs (L2) in light purple, and LINE/L2 in purple. **C,** Amino acid alignment of the partial PAZ domain of the PIWI1 protein among various Cyprinoidei fishes (red dot) and the outgroups (Nemacheilidae, blue dot). The phylogeny on the left panel represents the relationship among species. The orange cluster represents the Opsariichthyini tribe.

Drer: *Danio rerio*; Dcer: *Danionella cerebrum;* Lroh: *Labeo rohita*; Sgra: *Sinocyclocheilus graham*; Omac: *Onychostoma macrolepis*; Ptet: *Puntigrus tetrazona*; Ppro: *Pimephales promelas*; Agra: *Anabarilius graham*; Mmab: *Megalobrama amblycephala*; Cb: *Candidia barbatus*; Zp: *Zacco platypus;* Oe: *Opsariichthys evolans;* Op: *O. pachycephalus;* Ob: *O. bidens*; Ttib: *Triplophysa tibetana*; Tble: *T. bleekeri*

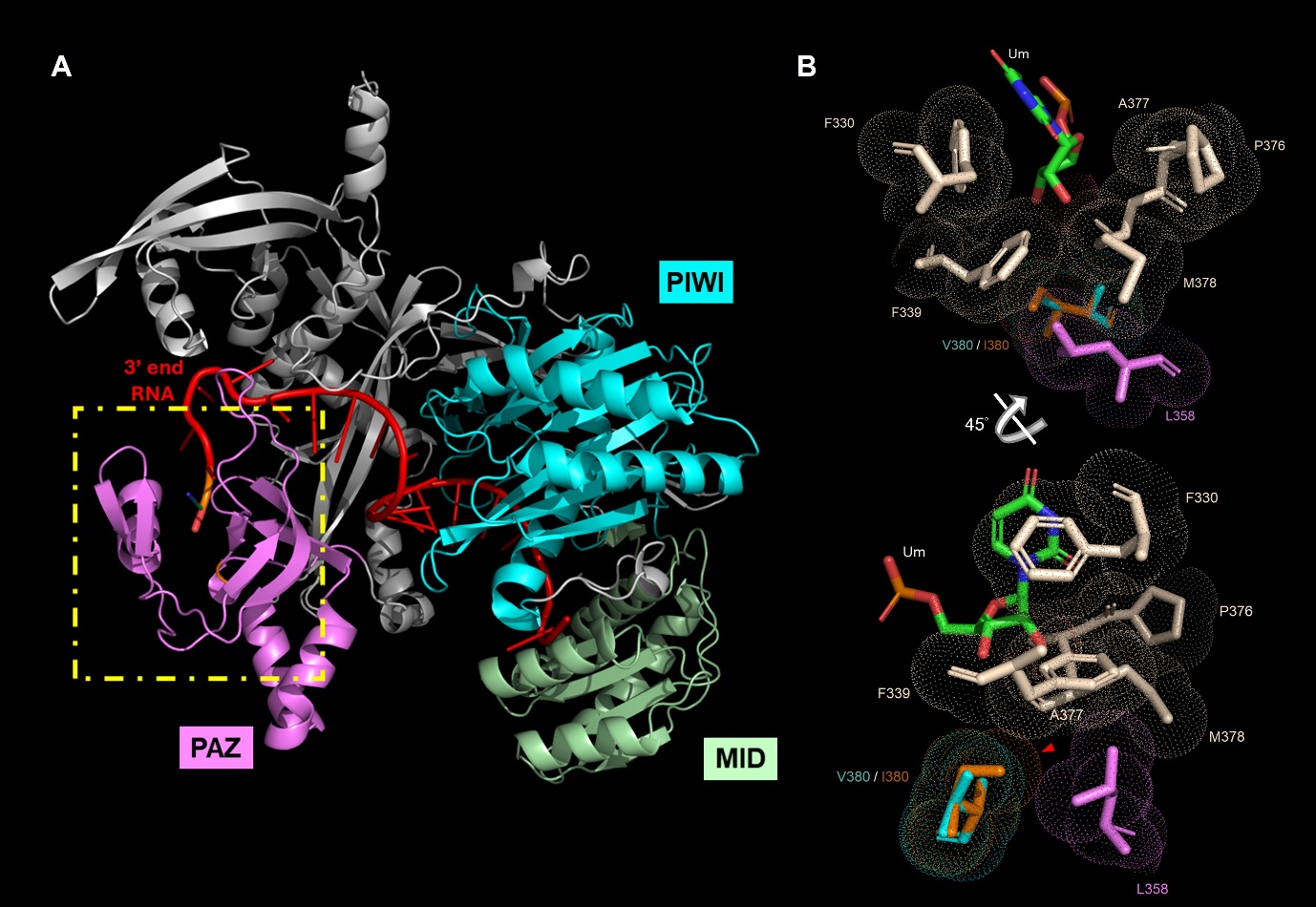

**Fig. S19.** **AlphaFold 3 predicted model of the PIWI-RNA complex.**

**A,** Overall structure of the PIWI-RNA complex with very low-confidence regions removed. The PAZ, MID, and PIWI domains are shown in violet, green, and cyan, respectively. The guide RNA (from PDB entry 3O7V) is shown in red. The yellow dashed box highlights the binding pocket, viewed from the same angle as in Figure S15B. **B,** Close-up of the binding pocket. Residues at the binding interface are shown in biscuit color. V380 and its mutant I380 are colored cyan and orange, respectively. The 380th residue is located beneath the binding interface (upper panel). An alternative view (lower panel) shows that the bulkier volume of I380 (red arrowhead) may shift adjacent residues (e.g., L358), thereby potentially alter the pocket stability. The dotted surface represents the van der Waals surface.

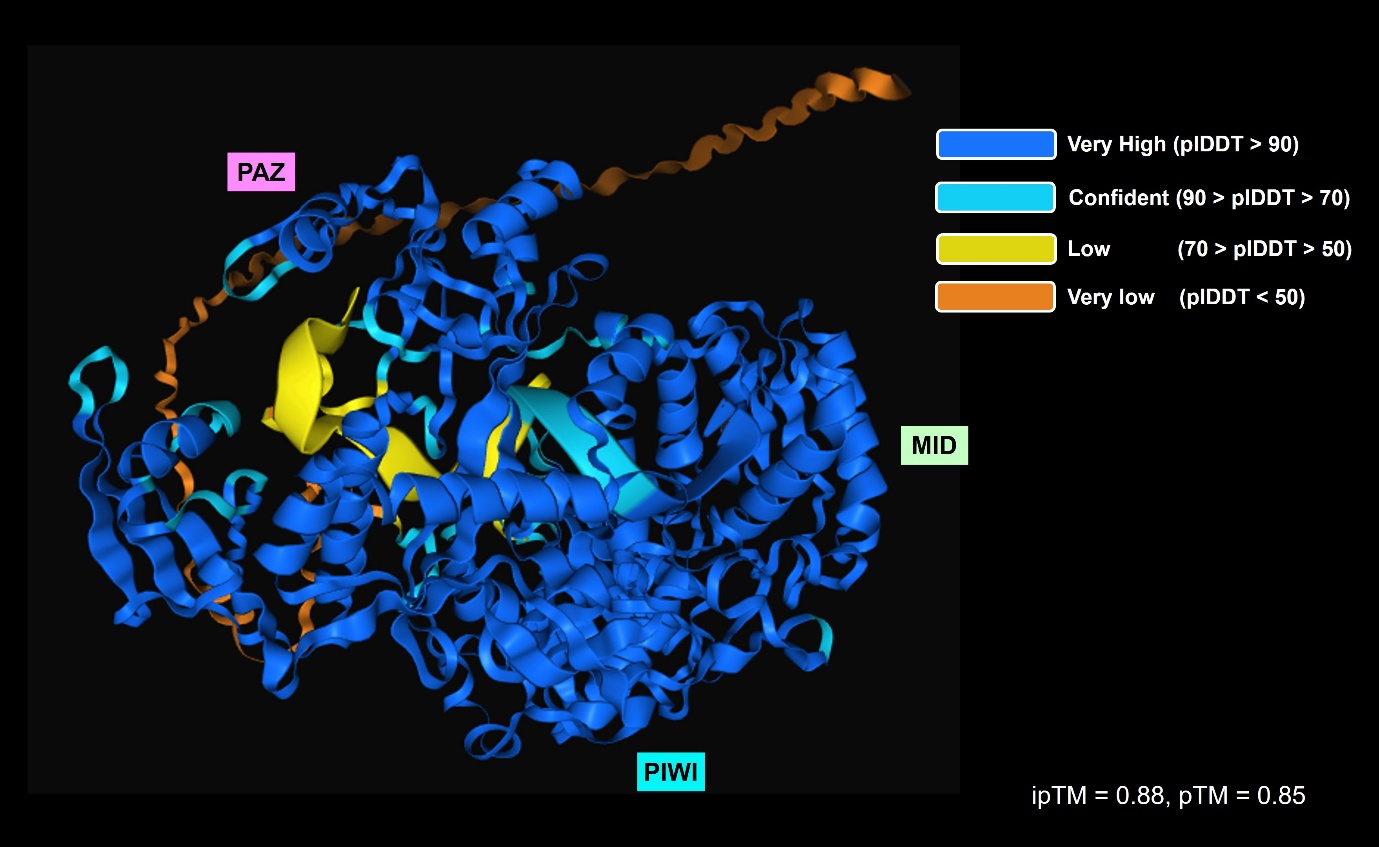

**Fig. S20.** **Confidence scores of the PIWI-RNA complex model.**

Regions with very high confidence are colored blue, confident regions are colored cyan, low-confidence regions are colored yellow, and very low-confidence regions are colored orange. pLDDT represents the per-residue confidence score, while ipTM denotes the interface predicted template modeling score, and pTM refers to the predicted template modeling score. For pTM, value above 0.5 means the overall prediction might be similar to the true structure. For iPTM, values higher than 0.8 represent a confident high-quality prediction.

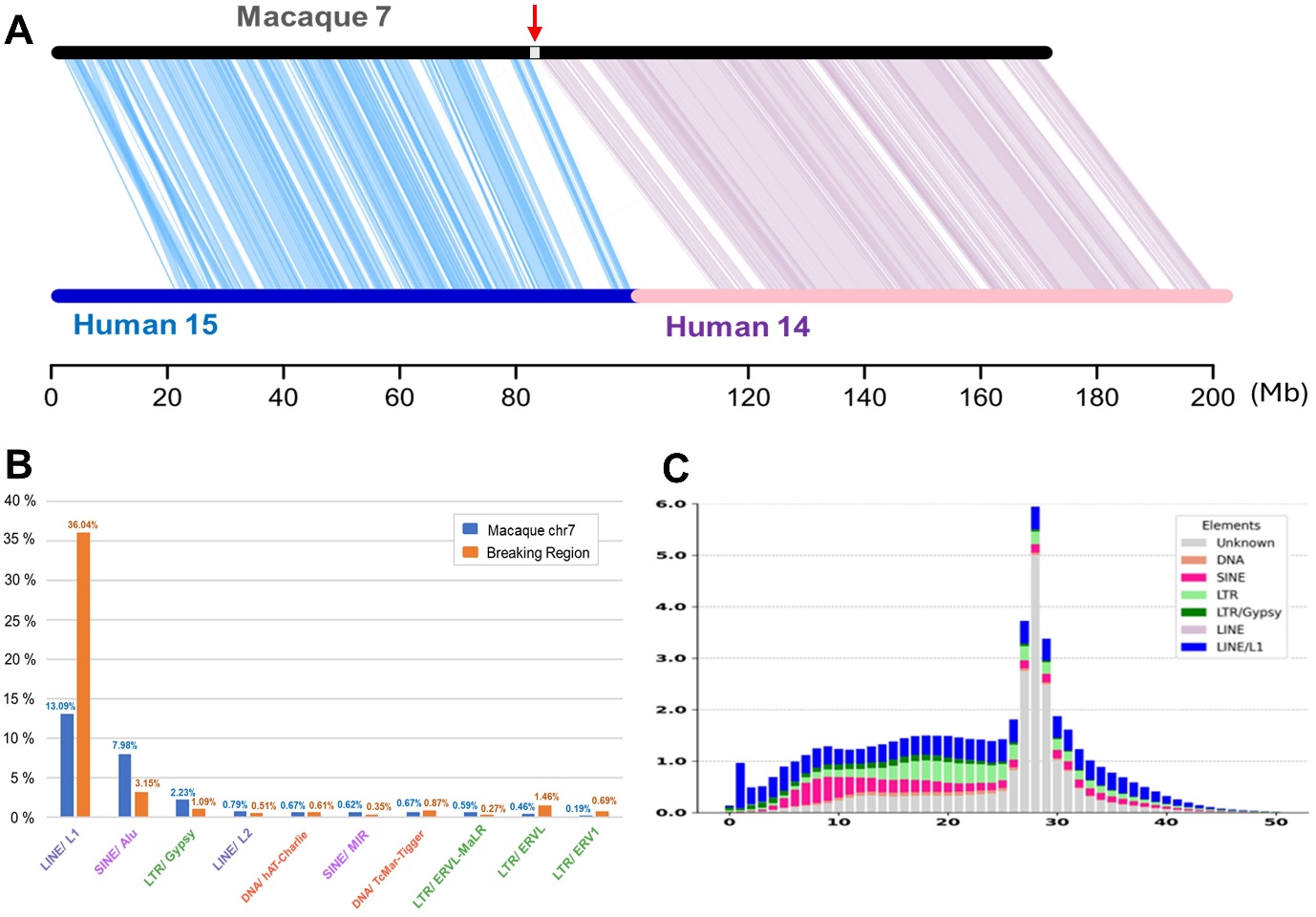

**Fig. S21.** **TEs in the breaking region Macaque chromosome 7.**

**A,** Syntenic relationships of Macaque chromosome 7 and two homologous fissioned chromosomes in human, chr14 and chr15. **B,** Proportion of different TEs in the Macaque chr7 and within breaking region. The proportion of L1 is three times higher in breaking regions than the chromosome average. **C,** The repeat landscape of Macaque genome. The reference genome of Macaque used in analysis is Mmul_10(GCF_003339765.1).

**Supplementary Tables**

**Table S1. Comparative karyotype of species used in this study.**

| Microscopic typing | | |  |  |  |  |  |  |
| --- | --- | --- | --- | --- | --- | --- | --- | --- |
| Species | Sex |  | Karyotype | | |  |  | Location* |
|  | Male | Female | N | M | SM | ST/T | FN |  |
| Cb | 1 | 1 | 24 | 9 | 12 | 3 | 45 | Xindian River, New Taipei |
| Zp | 2 | 1 | 24 | 9 | 11 | 4 | 44 | Xindian River, New Taipei |
| Oe | 2 | 2 | 39 | 2 | 1 | 36 | 42 | Xindian River, New Taipei |
| Op | 4 | 4 | 38 | 3 | 5 | 30 | 46 | Xindian River, New Taipei |
| Ob* | - | 1 | 39 | 2 | 2 | 35 | 43 | - |
| *In silico* typing | |  |  |  |  |  |  |  |
| Species | Sex |  | Karyotype | | |  |  |  |
|  | Male | Female | N | M | SM | ST/T | FN |  |
| Cb |  | 1 | 24 | 9 | 11 | 4 | 44 |  |
| Zp | 1 |  | 24 | 9 | 9 | 6 | 42 |  |
| Oe | 1 |  | 39 | 2 | 1 | 36 | 42 |  |
| Op |  | 1 | 38 | 2 | 3 | 33 | 43 |  |
| Ob^1^ |  | 1 | 39 | 1 | 1 | 37 | 41 |  |

1. Xu X et. al. 2024

*Sampling locations are shown in Fig. S1

N: haploid chromosome number; M: metacentric; SM: submetacentric; ST: sub-telocentric; T: telocentric; FN: functional number

**Table S2a. The statistic of sequence data used for genome assembly.**

| **Species** | **Cat. No.** | **Location*** | **Collected date** | **Accession number** | **Sample ID** | **Data Type** | **Sequencing Type** | **Library Size (Gb)** | **Number of reads** |
| --- | --- | --- | --- | --- | --- | --- | --- | --- | --- |
| *Candidia barbatus* | CbMS-B010 | Masu River, New Taipei | 2019.11.05 | *SRR30643960* | Cb_longReads | Genomic DNA | Oxford Nanopore Technology | 12.33 | 1,750,771 |
| *Candidia barbatus* | CbMS-B010 | Masu River, New Taipei | 2019.11.05 | *SRR30650513* | Cb_shortReads | Genomic DNA | Illumina pair-end 150 | 24.03 | 159,151,242 |
| *Candidia barbatus* | CbMS-B010 | Masu River, New Taipei | 2019.11.05 | *SRR30655453* | Cb_HiC | Genomic DNA | Hi-C | 69.48 | 463,178,016 |
| *Candidia barbatus* | CbMS-B010 | Masu River, New Taipei | 2019.11.05 | *SRR30650691* | Cb_eye | RNA-seq | Illumina pair-end 150 | 7.66 | 55,072,958 |
| *Candidia barbatus* | CbMS-B010 | Masu River, New Taipei | 2019.11.05 | *SRR30650704* | Cb_gill | RNA-seq | Illumina pair-end 150 | 7.50 | 55,280,462 |
| *Candidia barbatus* | CbMS-B010 | Masu River, New Taipei | 2019.11.05 | *SRR30650783* | Cb_liver | RNA-seq | Illumina pair-end 150 | 7.74 | 56,405,724 |
| *Candidia barbatus* | CbMS-B010 | Masu River, New Taipei | 2019.11.05 | *SRR30650804* | Cb_muscle | RNA-seq | Illumina pair-end 150 | 7.14 | 52,519,944 |
| *Candidia barbatus* | CbMS-B010 | Masu River, New Taipei | 2019.11.05 | *SRR30650803* | Cb_nose | RNA-seq | Illumina pair-end 150 | 7.59 | 55,935,306 |
| *Zacco platypus* | ZpHy926 | Keelung River, New Taipei | 2019.04.30 | *SRR30669803* | Zp_longReads | Genomic DNA | Oxford Nanopore Technology | 43.21 | 8,052,878 |
| *Zacco platypus* | ZpHy926 | Keelung River, New Taipei | 2019.04.30 | *SRR30669440* | Zp_shortReads | Genomic DNA | Illumina pair-end 150 | 28.59 | 189,345,098 |
| *Zacco platypus* | ZpZT041905 | Xindian River, New Taipei | 2023.04.19 | *SRR30669902* | Zp_HiC | Genomic DNA | Hi-C | 34.96 | 231,517,574 |
| *Zacco platypus* | ZpXD225M | Xindian River, New Taipei | 2020.08.06 | *SRR30669441* | Zp_225_eye | RNA-seq | Illumina pair-end 150 | 6.02 | 43,327,464 |
| *Zacco platypus* | Zp88 | Xindian River, New Taipei | 2018.03.31 | *SRR30669438* | Zp_brain | RNA-seq | Illumina pair-end 150 | 8.09 | 57,656,802 |
| *Zacco platypus* | Zp88 | Xindian River, New Taipei | 2018.03.31 | *SRR30669439* | Zp_nose | RNA-seq | Illumina pair-end 150 | 8.41 | 59,663,636 |
| *Zacco platypus* | Zp88 | Xindian River, New Taipei | 2018.03.31 | *SRR30669441* | Zp_eye | RNA-seq | Illumina pair-end 150 | 7.69 | 54,353,574 |
| *Opsariichthys evolans* | OeHMA602 | Houlong River, Miaoli | 2018.9.14 | *SRR31615236* | Oe_longReads | Genomic DNA | Oxford Nanopore Technology | 17.26 | 3,378,466 |
| *Opsariichthys evolans* | OeHMA602 | Houlong River, Miaoli | 2018.9.14 | *SRR31629970* | Oe_shortReads | Genomic DNA | Illumina pair-end 150 | 26.72 | 267,239,668 |
| *Opsariichthys evolans* | OeHS062505 | Sanxia River, New Taipei | 2023.06.25 | *SRR31629404* | Oe_HiC | Genomic DNA | Hi-C | 38.60 | 255,389,646 |
| *Opsariichthys evolans* | Oe29 | Houlong River, Miaoli | 2018.02.27 | *SRR31615677* | Oe_29_brain | RNA-seq | Illumina pair-end 150 | 8.72 | 60,896,714 |
| *Opsariichthys evolans* | Oe29 | Houlong River, Miaoli | 2018.02.27 | *SRR31628988* | Oe_29_eye | RNA-seq | Illumina pair-end 150 | 8.26 | 57,828,626 |
| *Opsariichthys evolans* | OeHMA602 | Houlong River, Miaoli | 2018.9.14 | *SRR31628990* | Oe_602_eye | RNA-seq | Illumina pair-end 150 | 7.70 | 53,677,072 |
| *Opsariichthys evolans* | Oe29 | Houlong River, Miaoli | 2018.02.27 | *SRR31628989* | Oe_29_nose | RNA-seq | Illumina pair-end 150 | 6.47 | 45,607,266 |
| *Opsariichthys evolans* | OeD239F | Houlong River, Miaoli | 2020.08.06 | *SRR31629087* | Oe_head | RNA-seq | Illumina pair-end 150 | 5.88 | 42,229,554 |
| *Opsariichthys pachycephalus* | OpHMA618 | Houlong River, Miaoli | 2018.09.29 | *SRR31656473* | Op_longReads | Genomic DNA | Oxford Nanopore Technology | 16.08 | 3,155,351 |
| *Opsariichthys pachycephalus* | OpMS062410 | Masu River, New Taipei | 2023.06.24 | *SRR31630025* | Op_shortReads | Genomic DNA | Illumina pair-end 150 | 9.76 | 97,552,680 |
| *Opsariichthys pachycephalus* | OpMS062406 | Masu River, New Taipei | 2023.06.24 | *SRR31630104* | Op_HiC | Genomic DNA | Hi-C | 37.50 | 248,250,294 |
| *Opsariichthys pachycephalus* | Op203 | Toucian River, Hsinchu | 2018.05.07 | *SRR31613698* | Op_brain | RNA-seq | Illumina pair-end 150 | 7.62 | 53,152,316 |
| *Opsariichthys pachycephalus* | Op203 | Toucian River, Hsinchu | 2018.05.07 | *SRR31613697* | Op_eye | RNA-seq | Illumina pair-end 150 | 7.28 | 51,193,382 |
| *Opsariichthys pachycephalus* | Op203 | Toucian River, Hsinchu | 2018.05.07 | *SRR31613696* | Op_nose | RNA-seq | Illumina pair-end 150 | 9.01 | 63,154,736 |
| *Opsariichthys bidens^1^* |  |  |  | *GWHBEIO00000000* |  | Genomic DNA |  |  |  |
| *Opsariichthys bidens^2^* |  |  |  | *SRR15969039* |  | RNA-seq | Illumina HiSeq pair-end 2500 | 6.94 | 46,241,418 |

1. Xu, X. et al (2022)

2. Ding, J. et al (2022)

*Sampling locations are shown in Fig. 20

**Table S2b. List of samples used for recombination rate analysis.**

| **Species** | **Cat. No.** | **Location*** | **Collected date** | **Accession number** | **Data Type** | **Sequencing Type** | **Instrument** | **Library Size (Gb)** | **Number of reads** | **Recombination rate** | **Geneflow** |
| --- | --- | --- | --- | --- | --- | --- | --- | --- | --- | --- | --- |
| *Candidia barbatus* | CbDP01 | Daping River, Hsinchu | 2023.06.25 | *SRR31673521* | Genomic DNA | Illumina pair-end 150 | Novaseq 6000 | 24.93 | 165,407,094 | O |  |
| *Candidia barbatus* | CbDP04 | Daping River, Hsinchu | 2023.10.29 | *SRR31674839* | Genomic DNA | Illumina pair-end 150 | Novaseq X plus | 9.26 | 61,327,740 | O |  |
| *Candidia barbatus* | CbMS-B010 | Masu River, New Taipei | 2019.11.05 | *SRR30650513* | Genomic DNA | Illumina pair-end 150 | HiSeq X | 24.03 | 159,151,242 | O |  |
| *Candidia barbatus* | CbMS02 | Masu River, New Taipei | 2023.06.24 | *SRR31704634* | Genomic DNA | Illumina pair-end 150 | Novaseq X plus | 10.41 | 68,963,192 | O |  |
| *Candidia barbatus* | CbWS01 | Waishuanghsi River, Taipei | 2023.10.07 | *SRR31705108* | Genomic DNA | Illumina pair-end 150 | Novaseq X plus | 9.34 | 61,879,196 | O |  |
| *Zacco platypus* | ZpHy926 | Keelung River, New Taipei | 2019.09.30 | *SRR30669440* | Genomic DNA | Illumina pair-end 150 | Novaseq 6000 | 28.59 | 189,345,098 | O | O |
| *Zacco platypus* | ZpRF310d | Keelung River, New Taipei | 2019.07.19 | *SRR31719979* | Genomic DNA | Illumina pair-end 150 | HiSeq X | 32.92 | 329,192,038 | O | O |
| *Zacco platypus* | ZpZT041901 | Xindian River, New Taipei | 2023.04.19 | *SRR31720049* | Genomic DNA | Illumina pair-end 150 | Novaseq 6000 | 17.19 | 114,086,130 | O |  |
| *Zacco platypus* | ZpZT041904 | Xindian River, New Taipei | 2023.04.19 | *SRR31720178* | Genomic DNA | Illumina pair-end 150 | Novaseq 6000 | 24.47 | 162,341,172 | O |  |
| *Zacco platypus* | ZpZT041905 | Xindian River, New Taipei | 2023.04.19 | *SRR31720186* | Genomic DNA | Illumina pair-end 150 | Novaseq 6000 | 21.02 | 139,482,516 | O |  |
| *Opsariichthys evolans* | OeHMA602 | Houlong River, Miaoli | 2018.09.14 | *SRR31629970* | Genomic DNA | Illumina pair-end 150 | Novaseq 6000 | 23.29 | 156,667,482 | O | O |
| *Opsariichthys evolans* | OeHS062501 | Sanxia River, New Taipei | 2023.06.25 | *SRR31824671* | Genomic DNA | Illumina pair-end 150 | Novaseq 6000 | 19.54 | 131,570,602 | O | O |
| *Opsariichthys evolans* | OeHS062507 | Sanxia River, New Taipei | 2023.06.25 | *SRR31824706* | Genomic DNA | Illumina pair-end 150 | Novaseq 6000 | 19.23 | 129,581,324 | O |  |
| *Opsariichthys evolans* | OEKL06 | Keelung River, New Taipei | 2020.08.02 | *SRR31824742* | Genomic DNA | Illumina pair-end 150 | Novaseq X plus | 11.17 | 76,313,464 | O |  |
| *Opsariichthys evolans* | OEKL07 | Keelung River, New Taipei | 2019.07.18 | *SRR31824898* | Genomic DNA | Illumina pair-end 150 | Novaseq X plus | 11.25 | 76,848,614 | O |  |
| *Opsariichthys evolans* | OEKL08 | Keelung River, New Taipei | 2019.07.18 | *SRR31824934* | Genomic DNA | Illumina pair-end 150 | Novaseq X plus | 11.53 | 79,073,056 | O |  |
| *Opsariichthys evolans* | OEWS01 | Waishuanghsi River, Taipei | 2024.01.07 | *SRR31825019* | Genomic DNA | Illumina pair-end 150 | Novaseq X plus | 10.84 | 74,378,028 | O |  |
| *Opsariichthys pachycephalus* | OPDA02 | Daan River, Miaoli | 2015.07.26 | *SRR31727918* | Genomic DNA | Illumina pair-end 150 | Novaseq X plus | 11.26 | 77,266,068 | O |  |
| *Opsariichthys pachycephalus* | OPDJ01 | Dajia River, Taichung | 2015.07.27 | *SRR31728056* | Genomic DNA | Illumina pair-end 150 | Novaseq X plus | 12.21 | 83,330,650 | O |  |
| *Opsariichthys pachycephalus* | OpDP062513 | Daping River, Hsinchu | 2023.06.25 | *SRR31824274* | Genomic DNA | Illumina pair-end 150 | Novaseq 6000 | 16.80 | 112,141,730 | O | O |
| *Opsariichthys pachycephalus* | OpHD337 | Houlong River, Miaoli | 2019.07.30 | *SRR31824275* | Genomic DNA | Illumina pair-end 150 | Novaseq 6000 | 17.46 | 116,666,228 | O | O |
| *Opsariichthys pachycephalus* | OpRF854 | Keelung River, New Taipei | 2019.07.23 | *SRR31824401* | Genomic DNA | Illumina pair-end 150 | HiSeq X | 12.19 | 83,326,768 | O |  |

*Sampling locations are shown in Fig. 20

**Table S3. Statistics of genome assembly.**

|  | ***Candidia barbatus*** | | ***Zacco platypus*** | | ***Opsariichthys pachycephalus*** | | ***Opsariichthys evolans*** | |
| --- | --- | --- | --- | --- | --- | --- | --- | --- |
|  | **Draft**  **assembly** | **HiC scaffolding** | **Draft**  **assembly** | **HiC scaffolding** | **Draft**  **assembly** | **HiC scaffolding** | **Draft**  **assembly** | **HiC scaffolding** |
| **Genome Size^a^** | 826.90 | 829.68 | 846.39 | 847.09 | 915.44 | 916.58 | 919.07 | 920.54 |
| **Number of contigs/scaffolds** | 1,739 | 1,233 | 2,543 | 2,332 | 2,749 | 2,483 | 2,931 | 2,835 |
| **Number of contigs/scaffolds > 1kbp** | 1,738 | 1,208 | 2,530 | 2295.00 | 2,742 | 2,420 | 2,928 | 2,776 |
| **N50^a^** | 2.70 | 32.96 | 6.17 | 31.42 | 1.54 | 21.86 | 0.95 | 21.75 |
| **L50** | 80 | 11 | 39 | 12 | 119 | 15 | 235 | 16 |
| **N90^a^** | 0.24 | 27.59 | 0.17 | 26.90 | 0.13 | 0.91 | 0.11 | 0.89 |
| **L90** | 456 | 22 | 379 | 23 | 976 | 37 | 1,356 | 38 |
| **Largest Scaffold/Contig Size^a^** | 15.73 | 46.66 | 18.26 | 46.34 | 20.51 | 42.83 | 9.35 | 40.09 |
| **Smallest Scaffold Size^a^** | NA | 23.02 | NA | 23.38 | NA | 8.07 | NA | 7.87 |
| **Depth** | - | 26x | - | 34x | - | 15x | - | 26x |
| **Mapping Rate^b^** | - | 99.32% | - | 99.74% | - | 99.82% | - | 99.31% |
| **BUSCO v5** | - | **97.97%** | - | **97.55%** | - | **97.69%** | - | **96.18%** |

a: In unit of Mb

b: Mapping short reads to the assembly genome

**Table S4. Genome statistics of the selected species.**

| **Species** | **Order** | **Chromosome Number** | **Genome Size** | **Contig Number** | **N50** | **L50** | **Busco v5** | **Source** | **Accession** |
| --- | --- | --- | --- | --- | --- | --- | --- | --- | --- |
| ***Candidia barbatus*** | **Xenocyprinidae** | **24** | **829.68 Mb** | **1,233** | **32.96 Mb** | **11** | **97.97%** | **This study** | **JBHGVQ000000000** |
| ***Zacco platypus*** | **Xenocyprinidae** | **24** | **846.39 Mb** | **2,332** | **31.42 Mb** | **12** | **97.55%** | **This study** | **JBHGZP000000000** |
| ***Opsariichthys pachycephalus*** | **Xenocyprinidae** | **38** | **916.58 Mb** | **2,483** | **21.86 Mb** | **15** | **97.69%** | **This study** | **JBJLZV000000000** |
| ***Opsariichthys evolans*** | **Xenocyprinidae** | **39** | **920.54 Mb** | **2,835** | **21.75 Mb** | **16** | **96.18%** | **This study** | **JBJLZW000000000** |
| *Opsariichthys bidens* (Female) | Xenocyprinidae | 39 | 818.78 Mb | 82 | 25.29 Mb | 13 | 97.30% | NGDC | GWHBEIO00000000 |
| *Megalobrama amblycephala* | Xenocyprinidae | 24 | 1109.89 Mb | 243 | 42.27 Mb | 11 | 97.50% | NCBI | GCA_018812025.1 |
| *Anabarilius grahami* | Xenocyprinidae | 24 | 991.89 Mb | 80,398 | 4.46 Mb | 60 | 97.20% | NCBI | GCA_003731715.1 |
| *Pimephales promelas* | Leuciscidae | 25 | 1066.43 Mb | 911 | 11.95 Mb | 23 | 96.80% | NCBI | GCF_016745375.1 |
| *Puntigrus tetrazona* | Cyprinidae | 25 | 730.82 Mb | 1,122 | 25.09 Mb | 13 | 97.80% | NCBI | GCF_018831695.1 |
| *Onychostoma macrolepis* | Cyprinidae | 25 | 886.57 Mb | 352 | 34.07 Mb | 11 | 97.30% | NCBI | GCA_012432095.1 |
| *Sinocyclocheilus grahami* | Cyprinidae | 25 | 1750.29 Mb | 31,277 | 1.16 Mb | 416 | 93.70% | NCBI | GCF_001515645.1 |
| *Labeo rohita* | Cyprinidae | 25 | 1126.54 Mb | 2,869 | 37.9 Mb | 12 | 98.20% | NCBI | GCA_022985175.1 |
| *Danionella cerebrum* | Danionidae | 25 | 735.3 Mb | 27,639 | 0.34 Mb | 604 | 90.40% | NCBI | GCA_007224835.1 |
| *Danio rerio* | Danionidae | 25 | 1679.2 Mb | 1,923 | 52.19 Mb | 14 | 95.90% | NCBI | GCF_000002035.6 |
| *Triplophysa tibetana* | Nemacheilidae | 25 | 652.93 Mb | 292 | 24.89 Mb | 11 | 96.60% | NCBI | GCA_008369825.1 |
| *Triplophysa bleekeri* | Nemacheilidae | 25 | 620.27 Mb | 181 | 22.89Mb | 12 | 92.90% | NCBI | VFQW00000000 |

**Table S5. List of percentage of repetitive sequence types.**

|  | **Total** | **DNA** | **LTR** | **LINE*** | **SINE** | **LINE/L2*** | **LTR/Gypsy*** |
| --- | --- | --- | --- | --- | --- | --- | --- |
| *Candidia barbatus* | 51.51% | 26.47% | 6.82% | 4.10% | 0.30% | 2.70% | 3.37% |
| *Zacco platypus* | 55.02% | 25.72% | 7.75% | 4.12% | 0.25% | 2.46% | 3.76% |
| *Opsariichthys evolans* | 53.20% | 27.06% | 7.48% | 3.81% | 0.30% | 2.22% | 3.20% |
| *Opsariichthys pachycephalus* | 55.38% | 26.80% | 8.25% | 4.70% | 0.28% | 2.91% | 3.52% |
| *Opsariichthys bidens^#^* | 51.63% | 26.69% | 7.95% | 4.29% | 0.31% | 2.70% | 3.86% |
| *Megalobrama amblycephala* | 62.18% | 32.68% | 9.42% | 3.56% | 0.20% | 1.79% | 4.22% |
| *Anabarilius grahami* | 37.70% | 30.45% | 5.52% | 3.62% | 0.49% | 1.73% | 2.58% |
| *Pimephales promelas* | 48.12% | 24.45% | 10.38% | 3.09% | 0.22% | 1.64% | 3.52% |
| *Puntigrus tetrazona* | 36.75% | 16.55% | 5.62% | 3.54% | 0.20% | 1.84% | 3.27% |
| *Onychostoma macrolepis* | 54.02% | 27.53% | 6.79% | 4.85% | 0.27% | 2.72% | 3.27% |
| *Sinocyclocheilus grahami* | 45.12% | 21.96% | 4.40% | 3.04% | 0.32% | 1.63% | 1.70% |
| *Labeo rohita* | 45.00% | 25.04% | 4.14% | 2.07% | 0.16% | 0.91% | 2.03% |
| *Danionella translucida* | 37.89% | 20.76% | 6.50% | 3.55% | 0.23% | 1.29% | 2.25% |
| *Danio rerio* | 64.45% | 46.19% | 8.57% | 4.44% | 1.03% | 1.90% | 2.69% |

*Significantly higher in the Opsariichthyini tribe (p < 0.05)

#Conducted by this study

**Table S6. Statistics of the predicted gene.**

| **Species** | **BUSCO v5** | **#gene** | **#mRNA** | **mean gene length** | **exons per mRNA** | **mean cds length(bp)** |
| --- | --- | --- | --- | --- | --- | --- |
| *Candidia barbatus* | 96.29% | 28,863 | 31,231 | 12,913 | 9.20 | 1,499 |
| *Zacco platypus* | 96.70% | 27,249 | 29,673 | 13,432 | 9.60 | 1,593 |
| *Opsariichthys pachycephalus* | 95.40% | 29,327 | 32,158 | 13,087 | 9.10 | 1,476 |
| *Opsariichthys evolans* | 91.10% | 30,730 | 33,910 | 12,002 | 8.60 | 1,372 |
| *Opsariichthys bidens** | 95.20% | 26,894 | 29,684 | 12,722 | 9.60 | 1,623 |
| *Opsariichthys bidens* | 93.50% | 23,992 | 23,992 | 16,469 | 9.80 | 1,670 |
| *Megalobrama amblycephala* | 99.42% | 29,877 | 59,927 | 22,873 | 13.30 | 2,161 |
| *Anabarilius grahami* | 62.20% | 23,906 | 23,906 | 9,022 | 7.20 | 1,414 |
| *Pimephales promelas* | 98.80% | 26,629 | 48,455 | 21,250 | 13.00 | 2,066 |
| *Puntigrus tetrazona* | 99.40% | 25,878 | 48,694 | 15,833 | 13.80 | 2,148 |
| *Onychostoma macrolepis* | 92.10% | 24,754 | 24,754 | 18,739 | 10.40 | 1,702 |
| *Sinocyclocheilus grahami* | 94.80% | 46,135 | 67,633 | 18,544 | 10.60 | 1,710 |
| *Labeo rohita* | 69.30% | 25,501 | 31,274 | 8,147 | 8.60 | 1,774 |
| *Danionella translucida* | 84.10% | 24,097 | 35,770 | 13,405 | 12.70 | 1,867 |
| *Danio rerio* | 98.80% | 32,749 | 57,108 | 30,798 | 13.00 | 2,113 |
| *Triplophysa tibetana* | 91.10% | 24,310 | 24,310 | 11,857 | 9.60 | 1,669 |

*Annotated by this study

**Table S7. Summary of chromosomal characteristics of Cb (Candidia barbatus) including TE enrichment in centromeres and breaking regions.**

|  |  |  |  | **Within centromere** | | | | **Within breaking region** | | | |  |  |  |  |
| --- | --- | --- | --- | --- | --- | --- | --- | --- | --- | --- | --- | --- | --- | --- | --- |
| **Chr** | **Centromere marker** | **Marker within centromere** | **Low recombination rate** | **L2 enrichment^1^** | **Gypsy enrichment^1^** | **L2 expansion^2^** | **Gypsy expansion^2^** | **L2 enrichment^1^** | **Gypsy enrichment^1^** | **L2 expansion^2^** | **Gypsy expansion^2^** | **Chromosome length*** | **centromere region*** | **breaking region start*** | **breaking region stop*** |
| 1 | O | O | O | O | O | O | O | O | O | O | O | 46.26 | 33.65 - 34.65 | 33.895 | 34.182 |
| 2 | O | O | O | O | O | O |  | O | O | O |  | 45.70 | 20.30 - 21.30 | 20.717 | 20.903 |
| 3 | O | O | O | O | O | O | O |  |  |  |  | 44.81 | 39.95 - 40.95 |  |  |
| 4 | O | O | O | O | O | O | O | O |  | O | O | 41.39 | 29.80 - 31.25 | 31.168 | 31.516 |
| 5 | O | O | O | O | O | O | O | O | O | O | O | 40.69 | 22.65 - 23.65 | 22.485 | 22.703 |
| 6 | O | O | O | O | O | O | O | O | O | O |  | 35.66 | 13.25 - 14.25 | 15.254 | 15.358 |
| 7 | O | O | O | O | O | O | O |  |  |  |  | 35.29 | 21.30 - 22.30 |  |  |
| 8 | O | O | O | O | O | O | O |  |  |  |  | 34.62 | 10.35 - 13.65 |  |  |
| 9 | O | O | O | O | O | O |  | O |  |  |  | 33.93 | 26.45 - 27.45 | 20.007 | 20.009 |
| 10 | O | O | O | O | O | O | O | O | O | O | O | 33.82 | 12.25 - 14.00 | 14.505 | 14.886 |
| 11 | O | O | O | O | O | O | O |  |  |  |  | 32.96 | 7.40 - 8.95 |  |  |
| 12 | O | O | O | O | O | O | O |  |  |  |  | 32.30 | 17.20 - 18.20 |  |  |
| 13 | O | O | O | O | O | O |  |  |  |  |  | 31.76 | 6.75 - 8.45 |  |  |
| 14 | O | O | O | O | O | O |  | O | O | O |  | 31.31 | 20.55 - 23.65 | 24.206 | 24.580 |
| 15 | O | O | O | O | O | O |  | O | O | O | O | 30.99 | 18.10 - 19.10 | 15.941 | 16.656 |
| 16 | O | O | O | O | O | O |  | O | O | O |  | 30.78 | 19.70 - 20.70 | 19.715 | 20.641 |
| 17 | O | O | O | O | O | O | O | O | O | O | O | 30.65 | 12.25 - 13.25 | 12.602 | 12.843 |
| 18 | O | O | O | O | O | O |  | O | O | O |  | 30.29 | 21.60 - 23.10 | 23.017 | 23.121 |
| 19 | O | O | O | O | O | O |  |  |  |  |  | 29.42 | 23.10 - 24.10 |  |  |
| 20 | O | O | O | O | O | O | O |  |  |  |  | 29.02 | 9.85 - 11.65 |  |  |
| 21 | O | O | O | O | O | O |  | O | O | O |  | 28.25 | 19.65 - 20.65 | 20.065 | 20.259 |
| 22 | O | O | O | O | O | O | O | O | O | O | O | 27.59 | 16.60 - 18.15 | 17.317 | 19.046 |
| 23 | O | O | O | O | O | O | O | O | O | O | O | 26.92 | 7.10 - 8.10 | 8.787 | 8.832 |
| 24 | O | O | O | O | O | O |  |  |  |  |  | 23.04 | 4.25 - 5.25 |  |  |

* In units of Mb.

1. L2 or Gypsy enrichments indicates an increased proportion of L2 or Gypsy

2. L2 or Gypsy expansion indicates a recent increase of L2 or Gypsy (K-distance < 5%)

**Table S8. Summary of chromosomal characteristics of Zp (Zacco platypus) including TE enrichment in centromere regions.**

|  |  |  |  | **Within centromere** | | | |  |  |  |  |
| --- | --- | --- | --- | --- | --- | --- | --- | --- | --- | --- | --- |
| **Chr** | **Centromere marker** | **Marker within centromere** | **Low recombination rate** | **L2 enrichment^1^** | **Gypsy enrichment^1^** | **L2 expansion^2^** | **Gypsy expansion^2^** | **Chromosome length*** | **Centromere region*** | **Breaking Region start*** | **Breaking Region stop*** |
| 1 | O | O | O | O | O | O | O | 46.17 | 12.70 - 13.70 | 12.976 | 13.348 |
| 2 | O | O | O | O | O | O | O | 44.61 | 23.45 - 24.45 | 23.844 | 23.975 |
| 3 | O | O | O | O | O | O | O | 41.49 | 30.10 - 31.10 | 30.395 | 30.765 |
| 4 | O | O | O | O | O | O | O | 40.37 | 38.30 - 39.30 |  |  |
| 5 | O | O | O | O | O | O | O | 40.04 | 21.10 - 22.10 | 20.594 | 21.618 |
| 6 | O | O | O | O | O | O | O | 36.38 | 31.35 - 32.35 |  |  |
| 7 | O | O | O | O | O | O | O | 35.12 | 14.95 - 15.95 | 14.963 | 15.511 |
| 8 | O | O | O | O | O | O | O | 35.00 | 9.45 - 10.45 | 9.796 | 10.297 |
| 9 | O | O | O | O | O | O | O | 33.23 | 19.80 - 20.80 | 18.810 | 21.112 |
| 10 | O | O | O | O | O | O | O | 32.86 | 13.55 - 14.55 | 13.609 | 13.899 |
| 11 | O | O | O | O | O | O | O | 31.51 | 11.65 - 12.65 |  |  |
| 12 | O | O | O | O | O | O | O | 31.42 | 14.20 - 15.20 | 14.124 | 15.632 |
| 13 | O | O | O | O | O | O | O | 31.41 | 7.10 - 8.70 |  |  |
| 14 | O | O | O | O | O | O | O | 31.23 | 14.75 - 16.65 |  |  |
| 15 | O | O | O | O | O | O | O | 30.38 | 11.30 - 13.05 | 11.315 | 11.640 |
| 16 | O | O | O | O | O | O | O | 30.07 | 6.80 - 7.80 | 7.177 | 7.346 |
| 17 | O | O | O | O | O | O | O | 30.02 | 6.85 - 7.85 |  |  |
| 18 | O | O | O | O | O | O | O | 29.51 | 17.60 - 18.60 | 18.036 | 18.081 |
| 19 | O | O | O | O | O | O | O | 27.87 | 4.90 - 5.90 |  |  |
| 20 | O | O | O | O | O | O | O | 27.72 | 7.65 - 8.65 | 7.986 | 8.267 |
| 21 | O | O | O | O | O | O | O | 27.26 | 16.90 - 17.90 | 16.929 | 17.785 |
| 22 | O | O | O | O | O | O | O | 26.99 | 17.85 - 18.85 | 17.271 | 18.800 |
| 23 | O | O | O | O | O | O | O | 26.90 | 18.05 - 19.05 |  |  |
| 24 | O | O | O | O | O | O | O | 23.42 | 18.15 - 19.15 |  |  |

* In units of Mb.

1. L2 or Gypsy enrichments indicates an increased proportion of L2 or Gypsy

2. L2 or Gypsy expansion indicates a recent increase of L2 or Gypsy (K-distance < 5%)

**Table S9. Summary of chromosomal characteristics of Oe (Opsariichthys evolans) including TE enrichment in centromere regions.**

|  |  |  |  |  | **Within centromere** | | | |  |  |
| --- | --- | --- | --- | --- | --- | --- | --- | --- | --- | --- |
| **Chr** | **Fission** | **Centromere marker** | **Marker within centromere** | **Low recombination rate** | **L2 enrichment^1^** | **Gypsy enrichment^1^** | **L2 expansion^2^** | **Gypsy expansion^2^** | **Chromosome length*** | **Putative centromere region*** |
| 1 |  | O | O | O | O | O | O | O | 40.09 | 0.00 - 1.00 |
| 2 | O |  |  | O | O | O | O | O | 36.68 | 0.00 - 1.00 |
| 3 |  | O | O | O | O | O | O |  | 36.03 | 0.00 - 1.00 |
| 4 |  | O | O | O | O | O | O | O | 35.70 | 20.90 - 25.15 |
| 5 |  | O | O | O | O | O | O | O | 33.97 | 25.55 - 26.55 |
| 6 |  | O |  | O | O | O | O | O | 33.46 | 14.50 - 15.50 |
| 7 | O |  |  | O | O | O | O |  | 31.69 | 30.69 - 31.69 |
| 8 |  | O | O |  | O | O | O | O | 31.06 | 5.90 - 6.90 |
| 9 |  | O | O | O | O | O | O | O | 30.56 | 0.60 - 1.60 |
| 10 | O | O | O | O | O | O | O | O | 26.64 | 25.64 - 26.64 |
| 11 | O | O | O | O | O | O | O | O | 26.03 | 25.03 - 26.03 |
| 12 | O |  |  | O | O | O | O | O | 25.07 | 0.00 - 1.00 |
| 13 |  |  |  | O | O | O | O | O | 24.49 | 23.49 - 24.49 |
| 14 |  | O | O | O | O | O | O |  | 24.35 | 23.35 - 24.35 |
| 15 | O | O | O | O | O | O | O |  | 22.55 | 21.55 - 22.55 |
| 16 | O |  |  | O | O | O | O | O | 21.75 | 0.00 - 1.00 |
| 17 | O | O | O | O | O | O | O | O | 21.61 | 14.45 - 17.15 |
| 18 | O | O |  | O | O | O | O |  | 21.32 | 20.32 - 21.32 |
| 19 | O | O | O | O | O | O | O | O | 21.19 | 20.19 - 21.19 |
| 20 | O |  |  | O | O | O | O |  | 21.08 | 20.08 - 21.08 |
| 21 | O | O | O | O | O | O | O | O | 20.69 | 18.90 - 19.90 |
| 22 | O | O | O | O | O | O | O | O | 20.00 | 0.00 - 1.00 |
| 23 | O |  |  | O | O | O | O | O | 20.00 | 19.00 - 20.00 |
| 24 | O | O | O | O | O | O | O | O | 19.08 | 0.00 - 1.55 |
| 25 | O | O | O | O | O | O | O | O | 18.91 | 0.00 - 1.00 |
| 26 | O | O | O | O | O | O | O | O | 17.81 | 0.00 - 1.00 |
| 27 | O |  |  | O | O | O | O | O | 15.72 | 0.00 - 1.00 |
| 28 | O | O | O | O | O | O | O | O | 14.55 | 13.55 - 14.55 |
| 29 | O |  |  | O | O | O | O | O | 14.42 | 13.42 - 14.42 |
| 30 | O | O |  | O | O | O | O | O | 14.15 | 0.00 - 1.00 |
| 31 | O |  |  | O | O | O | O | O | 13.10 | 0.00 - 1.00 |
| 32 | O | O |  | O | O | O | O | O | 13.05 | 12.05 - 13.05 |
| 33 | O | O | O | O | O | O | O |  | 12.52 | 0.00 - 1.60 |
| 34 | O | O | O | O | O | O | O | O | 12.35 | 0.00 - 1.00 |
| 35 | O |  |  | O | O |  | O |  | 10.72 | 0.00 - 1.00 |
| 36 | O | O | O | O | O | O | O | O | 10.48 | 9.48 - 10.48 |
| 37 | O | O | O | O | O | O | O | O | 9.99 | 7.70 - 8.70 |
| 38 | O | O | O | O | O | O | O | O | 8.87 | 0.00 - 1.00 |
| 39 | O | O |  | O | O | O | O | O | 7.87 | 0.00 - 1.00 |

* In units of Mb.

1. L2 or Gypsy enrichments indicates an increased proportion of L2 or Gypsy

2. L2 or Gypsy expansion indicates a recent increase of L2 or Gypsy (K-distance < 5%)

**Table S10. Summary of chromosomal characteristics of Op (Opsariichthys pachycephalus) including TE enrichment in centromere regions.**

|  |  |  |  |  | **Within centromere** | | | |  |  |
| --- | --- | --- | --- | --- | --- | --- | --- | --- | --- | --- |
| **Chr** | **Fission** | **Centromere marker** | **Marker within centromere** | **Low recombination rate** | **L2 enrichment^1^** | **Gypsy enrichment^1^** | **L2 expansion^2^** | **Gypsy expansion^2^** | **Chromosome length*** | **Putative centromere region*** |
| 1 |  | O | O | O | O | O | O | O | 42.83 | 40.00 - 41.00 |
| 2 | O | O | O | O | O | O | O | O | 37.06 | 0.00 - 2.30 |
| 3 |  | O | O | O | O | O | O | O | 37.06 | 12.75 - 14.80 |
| 4 |  | O | O | O | O | O | O | O | 36.52 | 32.65 - 33.65 |
| 5 |  | O | O | O | O | O | O | O | 33.82 | 7.65 - 9.40 |
| 6 |  | O | O | O | O | O | O | O | 33.73 | 17.05 - 18.05 |
| 7 |  | O | O | O | O | O | O | O | 32.71 | 0.60 - 1.60 |
| 8 | O |  |  | O | O | O | O | O | 32.00 | 0.00 - 1.00 |
| 9 |  | O | O | O | O | O | O | O | 31.92 | 7.50 - 8.50 |
| 10 |  | O | O | O | O | O | O | O | 30.79 | 5.80 - 8.45 |
| 11 | O | O | O | O | O | O | O | O | 27.34 | 26.34 - 27.34 |
| 12 | O | O | O | O | O | O | O | O | 26.90 | 0.00 - 1.30 |
| 13 |  |  |  | O | O | O | O | O | 25.36 | 3.50 - 4.50 |
| 14 |  | O | O | O | O | O | O | O | 24.57 | 0.00 - 1.00 |
| 15 | O | O | O | O | O | O | O | O | 21.86 | 20.15 - 21.15 |
| 16 | O | O | O | O | O | O | O | O | 21.76 | 0.05 - 1.05 |
| 17 | O | O | O | O | O | O | O | O | 21.75 | 0.00 - 1.00 |
| 18 | O | O | O | O | O | O | O | O | 21.74 | 0.10 - 1.10 |
| 19 | O | O | O | O | O | O | O | O | 20.30 | 0.00 - 1.00 |
| 20 | O | O | O | O | O | O | O | O | 20.02 | 19.02 - 20.02 |
| 21 | O | O | O | O | O | O | O | O | 19.88 | 18.88 - 19.88 |
| 22 | O | O |  | O | O | O | O | O | 19.62 | 0.00 - 1.00 |
| 23 | O |  |  | O | O | O | O | O | 19.54 | 0.00 - 1.00 |
| 24 | O | O | O | O | O | O | O | O | 18.72 | 18.22 - 19.22 |
| 25 | O | O | O | O | O | O | O | O | 18.72 | 17.72 - 18.72 |
| 26 | O | O |  | O | O | O | O | O | 17.32 | 0.00 - 1.00 |
| 27 | O | O | O | O | O | O | O | O | 14.49 | 13.49 - 14.49 |
| 28 | O |  |  | O | O | O | O | O | 14.46 | 0.00 - 1.00 |
| 29 | O | O | O | O | O | O | O | O | 13.88 | 12.88 - 13.88 |
| 30 | O | O |  | O | O | O | O | O | 13.55 | 0.00 - 1.00 |
| 31 | O | O | O | O | O | O | O | O | 12.35 | 11.05 - 12.05 |
| 32 | O | O | O |  | O | O | O | O | 12.16 | 9.45 - 10.45 |
| 33 | O |  |  | O | O | O | O | O | 11.98 | 10.98 - 11.98 |
| 34 | O | O |  | O | O | O | O | O | 11.37 | 10.37 - 11.37 |
| 35 | O | O |  | O | O | O | O | O | 10.15 | 0.00 - 1.00 |
| 36 | O | O | O | O | O | O | O | O | 9.19 | 8.19 - 9.19 |
| 37 | O | O | O | O | O | O | O | O | 9.13 | 0.00 - 1.00 |
| 38 | O | O | O | O | O | O | O | O | 8.07 | 0.00 - 1.00 |

* In units of Mb.

1. L2 or Gypsy enrichments indicates an increased proportion of L2 or Gypsy

2. L2 or Gypsy expansion indicates a recent increase of L2 or Gypsy (K-distance < 5%)

**Table S11. Summary of chromosomal characteristics of Ob (Opsariichthys bidens) including TE enrichment in centromere regions.**

|  |  |  |  |  | **Within centromere** | | | |  |
| --- | --- | --- | --- | --- | --- | --- | --- | --- | --- |
| **Chr** | **Fission** | **Cnetromere marker** | **marker within centromere** | **L2 enrichment^1^** | **Gypsy enrichment^1^** | **L2 expansion^2^** | **Gypsy expansion^2^** | **Chromosome length*** | **centromere region*** |
| 1 |  | O | O | O | O | O | O | 42.84 | 2.15 - 4.75 |
| 2 |  | O | O | O | O | O | O | 36.37 | 9.95 - 12.30 |
| 3 |  | O | O | O | O | O | O | 33.40 | 17.40 - 18.40 |
| 4 |  | O | O | O | O | O | O | 28.95 | 4.90 - 5.90 |
| 5 | O | O | O | O | O | O | O | 30.73 | 0.00 - 1.00 |
| 6 |  | O | O | O | O | O | O | 33.88 | 30.05 - 31.05 |
| 7 |  | O | O | O | O | O | O | 35.33 | 7.85 - 8.85 |
| 8 | O | O | O | O | O | O | O | 34.91 | 0.00 - 1.00 |
| 9 |  | O | O | O | O | O | O | 30.00 | 23.05 - 24.05 |
| 10 | O | O | O | O | O | O | O | 20.99 | 18.10 - 1.10 |
| 11 |  |  |  | O | O | O | O | 23.55 | 0.00 - 1.00 |
| 12 | O |  |  | O | O | O | O | 25.38 | 0.00 - 1.00 |
| 13 |  |  |  | O | O | O | O | 25.99 | 0.00 - 1.00 |
| 14 | O | O | O | O | O | O | O | 26.39 | 25.39 - 26.39 |
| 15 | O |  |  | O | O | O | O | 19.54 | 18.54 - 19.54 |
| 16 | O | O | O | O | O | O | O | 19.72 | 0.00 - 1.00 |
| 17 | O | O | O | O | O | O | O | 23.21 | 21.40 - 22.40 |
| 18 | O | O | O | O | O | O | O | 18.49 | 0.00 - 1.00 |
| 19 | O | O | O | O | O | O | O | 19.03 | 0.00 - 1.00 |
| 20 | O | O | O | O | O | O | O | 22.38 | 19.95 - 21.65 |
| 21 | O | O | O | O | O | O | O | 20.70 | 19.20 - 20.20 |
| 22 | O | O | O | O | O | O | O | 20.82 | 0.00 - 1.00 |
| 23 | O | O | O | O | O | O | O | 25.29 | 24.29 - 25.29 |
| 24 | O | O | O | O | O | O | O | 20.92 | 0.00 - 1.00 |
| 25 | O | O | O | O | O | O | O | 18.93 | 17.93 - 18.93 |
| 26 | O | O |  | O | O | O | O | 16.96 | 15.96 - 16.96 |
| 27 | O | O | O | O | O | O | O | 12.35 | 0.00 - 1.00 |
| 28 | O | O | O | O | O | O | O | 15.25 | 0.10 - 1.10 |
| 29 | O | O | O | O | O | O | O | 12.86 | 11.86 - 12.86 |
| 30 | O | O | O | O | O | O | O | 9.89 | 0.00 - 1.00 |
| 31 | O | O | O | O | O | O | O | 11.31 | 0.00 - 1.00 |
| 32 | O | O | O | O | O | O | O | 11.07 | 9.65 - 11.07 |
| 33 | O | O | O | O | O | O | O | 13.13 | 11.85 - 12.85 |
| 34 | O | O |  | O | O | O | O | 10.66 | 0.00 - 1.00 |
| 35 | O | O |  | O | O | O | O | 12.29 | 0.00 - 1.00 |
| 36 | O | O | O | O | O | O | O | 8.80 | 0.00 - 1.00 |
| 37 | O | O | O | O | O | O |  | 6.85 | 0.00 - 1.00 |
| 38 | O | O |  | O | O | O | O | 6.77 | 5.77 - 6.77 |
| 39 | O | O | O |  |  | O |  | 8.80 | 7.80 - 8.80 |

* In units of Mb.

1. L2 or Gypsy enrichments indicates an increased proportion of L2 or Gypsy

2. L2 or Gypsy expansion indicates a recent increase of L2 or Gypsy (K-distance < 5%)

**Table S12. lnCL of each model.**

|  | ***Zacco platypus*** versus | | | |
| --- | --- | --- | --- | --- |
|  | ***O. evolans*** | *△lnCL* | ***O. pachycephalus*** | *△lnCL* |
| **DIV** | -93,217,569 | -4,959 | -95,610,326 | -1,664 |
| **IM_->Zacco_** | -93,212,610 | 0 | -95,608,662 | 0 |
| **IM_->Opsariichthys_** | -93,212,627 | -17 | -95,608,832 | -170 |
| **MIG_->Zacco_** | -95,188,968 | -1,976,358 | -97,817,329 | -2,208,667 |
| **MIG_->Opsariichthys_** | -95,448,339 | -2,235,729 | -97,786,873 | -2,178,211 |

**Table S13. Chromosomal homology across species of opsariichthyin.**

| ***Candidia barbatus*  (Cb)** | ***Zacco platypus*  (Zp)** | ***Opsariichthys pachycephalus* (Op)** | ***Opsariichthys evolans*  (Oe)** | ***Opsariichthys bidens*  (Ob)** |
| --- | --- | --- | --- | --- |
| Cb1(R) | Zp1 | Op30(R) | Oe30(R) | Ob35(R) |
|  |  | Op2 | Oe2 | Ob8 |
| Cb2(R) | Zp2 | Op12 | Oe11(R) | Ob23(R) |
|  |  | Op18 | Oe16 | Ob17(R) |
| Cb4 | Zp3 | Op8(R) | Oe7 | Ob5(R) |
|  |  | Op34(R) | Oe34 | Ob34 |
| Cb3 | Zp4 | Op1 | Oe1(R) | Ob1(R) |
| Cb5 | Zp5 | Op17(R) | Oe17 | Ob20 |
|  |  | Op22 | Oe23(R) | Ob10(R) |
| Cb7 | Zp6 | Op4 | Oe3(R) | Ob6 |
| Cb6 | Zp7 | Op28(R) | Oe27(R) | Ob28(R) |
|  |  | Op19 | Oe18(R) | Ob22 |
| Cb14(R) | Zp8 | Op38(R) | Oe38(R) | Ob37(R) |
|  |  | Op11(R) | Oe10(R) | Ob14(R) |
| Cb9 | Zp9 | Op15 | Oe20 | Ob25 |
|  |  | Op31(R) | Oe32(R) | Ob33(R) |
| Cb10 | Zp10 | Op27 | Oe29 | Ob29 |
|  |  | Op21(R) | Oe19(R) | Ob15(R) |
| Cb8 | Zp11 | Op3 | Oe4(R) | Ob2 |
| Cb15(R) | Zp12 | Op29 | Oe28 | Ob27(R) |
|  |  | Op26 | Oe26 | Ob26(R) |
| Cb11 | Zp13 | Op5 | Oe5(R) | Ob7 |
| Cb12 | Zp14 | Op6 | Oe6(R) | Ob3 |
| Cb16(R) | Zp15 | Op33 | Oe31(R) | Ob31(R) |
|  |  | Op20(R) | Oe21(R) | Ob16 |
| Cb18(R) | Zp16 | Op9 | Oe39(R) | Ob38 |
|  |  |  | Oe12 | Ob12 |
| Cb13 | Zp17 | Op7 | Oe8 | Ob9(R) |
| Cb17(R) | Zp18 | Op25 | Oe25(R) | Ob21 |
|  |  | Op32(R) | Oe33 | Ob32(R) |
| Cb19(R) | Zp19 | Op10 | Oe9 | Ob4 |
| Cb21(R) | Zp20 | Op37(R) | Oe36 | Ob39 |
|  |  | Op16(R) | Oe15 | Ob24(R) |
| Cb22 | Zp21 | Op24 | Oe24(R) | Ob18(R) |
|  |  | Op35 | Oe35 | Ob30 |
| Cb23(R) | Zp22 | Op23(R) | Oe22(R) | Ob19(R) |
|  |  | Op36(R) | Oe37(R) | Ob36 |
| Cb20(R) | Zp23 | Op13(R) | Oe14 | Ob13(R) |
| Cb24(R) | Zp24 | Op14 | Oe13(R) | Ob11 |

The orientation of chromosome sequences is determined based on Zp, with 'R' indicating a reversed orientation compared to Zp.
